# Feeding habits and novel prey of larval fishes in the northern San Francisco Estuary

**DOI:** 10.1101/2020.10.18.344440

**Authors:** Michelle J. Jungbluth, Jillian Burns, Lenny Grimaldo, Anne Slaughter, Aspen Katla, Wim Kimmerer

## Abstract

Food limitation can dampen survival and growth of fish during early development. To investigate prey diversity important to the planktivorous larval longfin smelt (*Spirinchus thaleichthys*) and Pacific herring (*Clupea pallasii*) from the San Francisco Estuary, we used DNA metabarcoding analysis of the cytochrome oxidase I gene on the guts of these fishes and on environmental zooplankton samples. Differential abundance analysis suggested that both species consumed the most abundant zooplankton at a lower rate than their availability in the environment. Both fish consumed the prey that were commonly available and relatively abundant. Prey taxa substantially overlapped between the two species (Schoener’s index = 0.66), and alpha diversity analysis suggested high variability in the content of individual guts. Abundant prey taxa in both fish species included the copepods *Eurytemora carolleeae, Acanthocyclops americanus*, and *A. robustus*; the *Acanthocyclops* spp. are difficult to identify morphologically. A few uncommon prey in the diets hint at variable feeding strategies, such as herring (presumably egg) DNA in the longfin smelt diets, which suggests feeding near substrates. Herring consumed the small (<0.5 mm) copepod *Limnoithona tetraspina* more frequently (30%) than did smelt (2%), possibly indicating differences in foraging behavior or sensory abilities. Among the unexpected prey found in the diets was the cnidarian *Hydra oligactis*, the polychaete *Dasybranchus* sp., and a newly identified species *Mesocyclops pehpeiensis*. “Unknown” DNA was in 56% of longfin smelt diets and 57% of herring diets, and made up 17% and 21% of the relative read abundance in the two species, respectively. Our results suggest that these two fishes, which overlap in nursery habitat, also largely overlap in food resources necessary for larval survival.

## Introduction

Estuaries provide critical nursery habitats for many coastal and anadromous species including forage fishes (Beck *et al*., 2001; Boehlert & Mundy 1988), which are an important link between plankton and higher-level predators (Cury *et al*., 2000; Cury *et al*., 2011; Hunt *et al*., 2002). Knowledge of the prey that support recruitment of forage fishes in estuarine nursery habitats could improve management of these habitats. While some studies have investigated the diets of larval fishes in estuarine habitats, few have applied the high level of taxonomic resolution available through dietary DNA (dDNA) metabarcoding and, as a result, are likely to miss many potentially important or informative taxa. Larval fishes feed on small planktonic organisms that can be hard to identify visually, and some softer-bodied organisms may be digested more rapidly, limiting the resolution of traditional morphological diet analysis and thus limiting our knowledge of the breadth of prey important to fish recruitment (Llopiz 2013; Montagnes *et al*., 2010).

In the northern, brackish to fresh reaches of the San Francisco Estuary (SFE), key forage fish species of management interest have declined substantially over the last few decades (Sommer et al., 2007), including longfin smelt *Spirinchus thaleichthys* (state-listed as threatened), delta smelt (*Hypomesus transpacificus*; state and federally listed) and striped bass *Morone saxatilis*. A downward shift in overall prey abundance and a change in prey availability are both implicated as major factors responsible for the declining trend of forage fish in the low salinity zone of the SFE (Kimmerer 2002; Kimmerer 2006; Kimmerer & Orsi 1996; Orsi & Ohtsuka 1999; Sommer et al., 2007; Thompson et al., 2010; Rose et al., 2013). The reduction in prey abundance is mostly attributable to grazing effects caused by the non-native Asian clam *Potamacorbula amurensis* following its establishment in the estuary in the late 1980’s. Prey availability in the low-salinity zone shifted from a zooplankton community numerically dominated by calanoid copepods to one dominated by a small cyclopoid copepod, *Limnoithona tetraspina* (Bouley & Kimmerer 2006).

Growth and survival of larval fish in estuarine ecosystems is strongly linked to suitable prey availability and associated energetic costs of capturing prey while maintaining position in a tidal environment (Leggett & Deblois 1994; Pepin 2004; Pepin *et al*., 2014). Thus, both declines in prey abundance and shifts in prey availability could affect the successful recruitment of forage fish larvae in the northern SFE. Diet shifts have been documented for juvenile and adult fish in the SFE that have undergone significant declines in abundance, including striped bass and longfin smelt (Bryant & Arnold 2007; Feyrer et al., 2003). In other cases, the decline in prey abundance has caused some forage fish in the northern SFE to shift their distribution seaward (Kimmerer 2006) or towards different habitats where foraging opportunities may be better (Sommer et al., 2011).

The impact of the food web collapse in the northern SFE to larval life stages of forage fish is less understood but may be important for understanding interannual variation in recruitment. Here, we study the diet patterns of larval longfin smelt and Pacific herring. Longfin smelt are small pelagic fish that use the lower salinity habitat in the SFE for spawning and larval rearing, while sub-adults and adults are thought to rear primarily in the ocean or San Francisco Bay. Both the distribution and abundance of longfin smelt varies strongly with freshwater flow. During years of higher freshwater flow into the estuary, larval longfin smelt rearing shifts seaward and into shallow marsh habitats (Grimaldo et al., 2017; 2020). Correspondingly, age-0 longfin smelt abundance is higher by almost two orders of magnitude during years with higher freshwater flow compared to years when freshwater flow is low (Kimmerer et al., 2009). Rearing conditions may be enhanced during high-flow years but mechanisms for effects of flow on rearing conditions are still largely unknown (Grimaldo et al., 2020). Pacific herring spawn adhesive eggs on substrates mainly in the seaward regions of the estuary but larvae can be broadly distributed throughout the estuary depending on freshwater flow. For example, Grimaldo et al., (2020) observed larval herring in relatively modest abundance in the northern estuary during a low flow year which may be due either net landward movement via two-layer gravitational flow or some local spawning activity.

Rearing of longfin smelt and Pacific herring larvae may overlap in the SFE in some years (Grimaldo et al., 2020), thus we would expect them to have similar diets through foraging in a shared prey field when their spatial distributions are similar. Studies of feeding by larval longfin smelt to date are rare, but morphological gut content analyses of larvae have found copepods to be important prey, including *Acanthocyclops* spp., *Pseudodiaptomus forbesi* (Hobbs *et al*., 2006), and *Eurytemora carolleeae* (formerly *E. affinis*; Alekseev & Souissi 2011; Slater 2015). Herring diet studies in Central San Francisco Bay (7-9 mm larvae) found tintinnid ciliates to be the most common prey, followed by juvenile copepods, diatoms, gastropod veligers, and unknown crustaceans (Bollens & Sanders 2004). In other studies, larval Pacific herring consumed calanoid and harpacticoid copepods (Bowers & Williamson 1951; Wailes 1936), diatoms, rotifers (Choi *et al*., 2015), cirripede and gastropod larvae (Blaxter 1965; Bowers & Williamson 1951; Wailes 1936), fish eggs, and Artemia sp. (Blaxter 1965; Kurata 1959). To date there has been no direct comparison of prey consumed by larval longfin smelt and Pacific herring when their distributions overlap.

Morphological gut content analysis is the most direct method for analyzing feeding and has been a mainstay of aquatic trophic studies for over a century (Evermann & Lee 1906). Like any single method, this type of analysis has several limitations: identification of prey depends on the skill of the analyst, guts may contain unidentifiable material or appear empty but still contain prey remains (e.g., Slater & Baxter 2014), and variable digestion time of the prey can bias results toward hard-bodied prey (Hyslop 1980). All of these limitations can lead to an incomplete understanding of aquatic food webs (Pompanon *et al*., 2012; Sousa *et al*., 2019). Although arthropods are important prey for larval fishes (Llopiz 2013), many soft-bodied plankton can provide important nutrition to larval fishes. For example, protists may alleviate food limitation in larval fish (Hunt von Herbing & Gallager, 2000; Stoecker & Capuzzo 1990) and may be assimilated more easily than invertebrates (Reitan *et al*., 1998).

Application of DNA metabarcoding to investigate diets has successfully identified a wide range of previously unknown prey in a wide variety of aquatic species (Pompanon *et al*., 2012; Roslin & Majaneva 2016; Sousa *et al*., 2019) including lobster larvae (O’Rorke *et al*., 2012), copepods (Harfman *et al*., 2019; Ho *et al*., 2017), copepod nauplii (Craig *et al*., 2014), and fishes (e.g., Albaina *et al*., 2016; Hirai *et al*., 2017; Leray *et al*., 2019; Waraniak et al, 2019). This is possible because molecular techniques can detect and identify specific prey DNA in a predator’s digestive system down to a few copies of a gene, long after the prey’s body has decomposed beyond morphological recognition (Pompanon *et al*., 2012; Sousa *et al*., 2019). While no dDNA studies have been done on the larvae of Pacific herring or longfin smelt to date, dDNA studies of other clupeid larvae and juveniles revealed diverse prey including numerous species of copepods, decapods, ostracods, cnidarians, as well as phyllodocid and capitellid polychaetes, echinoderms, bivalves, gastropods and other fish species (Bowser *et al*., 2013; Hirai *et al*., 2017). These results suggest that Pacific herring and longfin smelt likely consume more diverse prey taxa than has been detected by morphological diet analysis.

We identified the prey in larval longfin smelt and Pacific herring guts collected during the same time period in the SFE to determine diet similarity based on an ambient prey field. We applied metabarcoding analysis of the mitochondrial cytochrome oxidase subunit I (mtCOI) gene across diverse taxa to detect and identify common prey taxa in guts that may be missed by traditional morphological methods. Feeding specialization among habitats was inferred for the two species from the diet analysis.

## Materials and Methods

### Study Area and Sample Collection

The San Francisco Estuary, the largest estuary on the west coast of the United States,which drains about 40 percent of California’s area. The climate is mediterranean with most of the freshwater runoff occurring in winter to early spring, and most flows through the California Delta linking the Sacramento and San Joaquin rivers through Suisun, San Pablo, and San Francisco Bays to the Pacific Ocean (Fig. 1). Other sources of freshwater in the estuary include small streams and large tributaries (e.g., the Petaluma River). Fish and zooplankton were collected for this study during the spring of 2017, which was the year of highest freshwater flow from 1955 to 2020 (Grimaldo et al., 2020). During high flow years, distributions of pelagic fishes such as longfin smelt and Pacific herring shift seaward, generally tracking the salinity distribution (Grimaldo et al., 2020).

**Figure 1:**
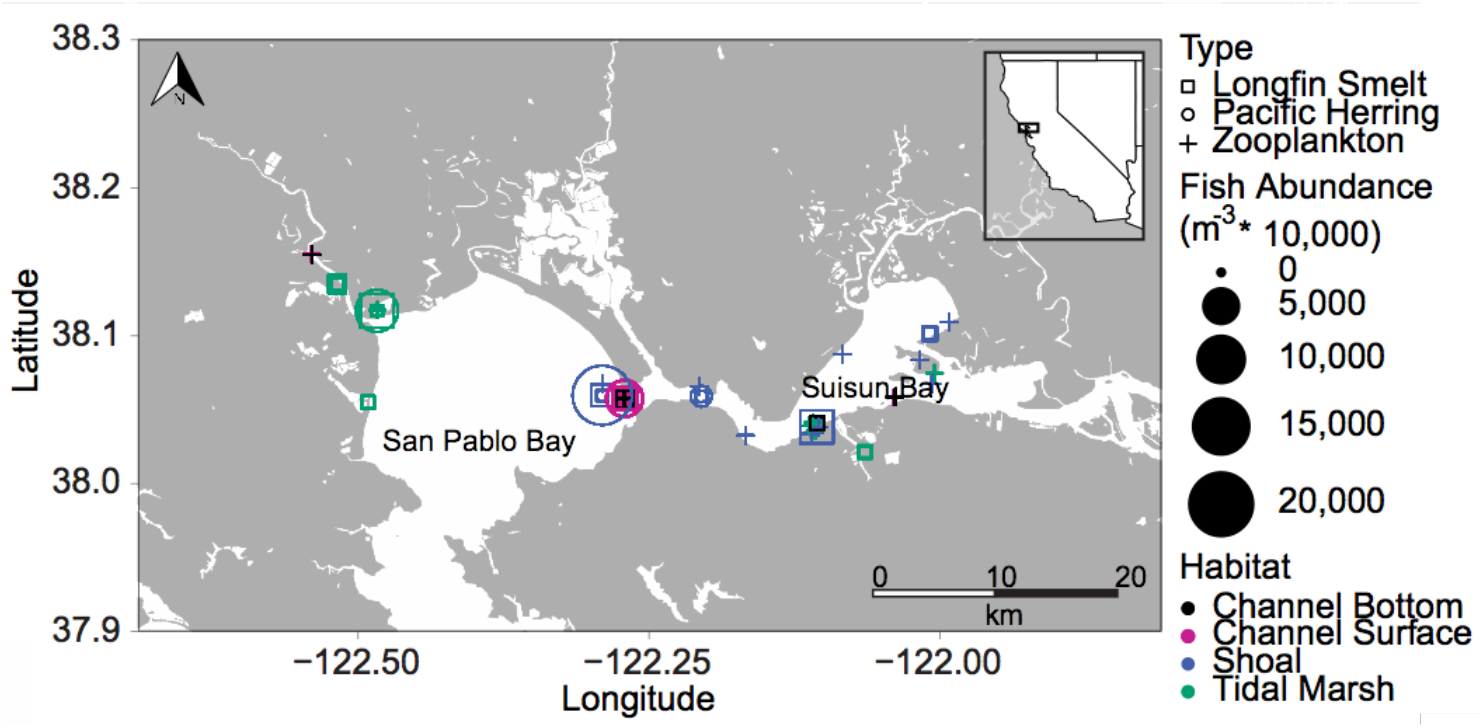
Map of sample sites in the northern San Francisco Estuary; inset shows location in California, western United States. Sites were surveyed for longfin smelt and Pacific herring. Shapes indicate where longfin smelt (squares), Pacific herring (circles), or zooplankton without fish (cross) were sampled. Zooplankton were also sampled where fish were collected. For squares and circles, the size of the point indicates the fish abundance (m^−3^ × 10,000). Habitats are indicated with color: channel bottom (black), channel surface (pink), shoal (blue), or tidal marsh (green).

Larval fish and zooplankton were collected across different habitats in San Pablo Bay and Suisun Bay of the northern SFE (Figure 1, Table 1). San Pablo Bay and Suisun Bay are shallow brackish embayments connected by a narrow deep channel (Carquinez Strait). Sampling occurred between 28 February 2017 and 25 May 25 2017 for a total of 32 samples. Sites were identified based on habitat and location and selected for sampling based on a randomly stratified sample design. The four habitat types sampled in both regions comprised shallow open water shoals, channels of tidal marshes, and deep open channels near the water surface (channel surface) and near the bottom (channel bottom).

**Table 1.** Sample information corresponding to tow numbers where zooplankton and fish samples were collected.

| Date | Time | Tow # | Latitude | Longitude | Region | Habitat | Fish Present |
| --- | --- | --- | --- | --- | --- | --- | --- |
| 2/28/17 | 09:22 | 1 | 38.0674 | -122.2900 | San Pablo | Shoal | - |
| 2/28/17 | 09:46 | 2 | 38.0574 | -122.2714 | San Pablo | Channel Surface | - |
| 2/28/17 | 10:11 | 3 | 38.0574 | -122.2714 | San Pablo | Channel Bottom | - |
| 2/28/17 | 11:17 | 4 | 38.0338 | -122.1105 | San Pablo | Shoal | - |
| 2/28/17 | 11:52 | 5 | 38.0327 | -122.1670 | San Pablo | Shoal | - |
| 3/7/17 | 08:13 | 6 | 38.0368 | -122.1044 | Suisun | Tidal Marsh | F |
| 3/7/17 | 08:53 | 7 | 38.0384 | -122.1039 | Suisun | Shoal | - |
| 3/7/17 | 09:46 | 8 | 38.0674 | -122.0068 | Suisun | Shoal | - |
| 3/7/17 | 10:51 | 9 | 38.0837 | -122.0171 | Suisun | Shoal | - |
| 3/7/17 | 11:58 | 10 | 38.0875 | -122.0838 | Suisun | Shoal | - |
| 3/7/17 | 12:49 | 11 | 38.0583 | -122.0391 | Suisun | Channel Surface | - |
| 3/7/17 | 13:32 | 12+ | 38.0586 | -122.0389 | Suisun | Channel Bottom | - |
| 3/9/17 | 08:32 | 13+ | 38.0576 | -122.2709 | San Pablo | Channel Bottom | F |
| 3/9/17 | 09:54 | 14 | 38.0572 | -122.2705 | San Pablo | Channel Bottom | F |
| 3/9/17 | 11:34 | 15+ | 38.1170 | -122.4838 | San Pablo | Tidal Marsh | F |
| 3/9/17 | 12:44 | 16+ | 38.1350 | -122.5181 | San Pablo | Channel Surface | F |
| 3/9/17 | 13:37 | 17 | 38.1552 | -122.5401 | San Pablo | Channel Surface | F |
| 3/9/17 | 14:22 | 18 | 38.1551 | -122.5399 | San Pablo | Channel Bottom | - |
| 3/22/17 | 08:51 | 19 | 38.0359 | -122.1071 | Suisun | Tidal Marsh | - |
| 3/22/17 | 09:30 | 20+ | 38.0385 | -122.1058 | Suisun | Shoal | F |
| 3/22/17 | 10:41 | 21 | 38.0411 | -122.1056 | Suisun | Channel Bottom | F |
| 3/22/17 | 13:03 | 22 | 38.1015 | -122.0085 | Suisun | Shoal | F |
| 3/23/17 | 07:26 | 23+ | 38.0591 | -122.2050 | Suisun | Shoal | F |
| 3/23/17 | 08:37 | 24 | 38.0598 | -122.2902 | San Pablo | Shoal | F |
| 3/23/17 | 09:37 | 25 | 38.1175 | -122.4835 | San Pablo | Tidal Marsh | F |
| 3/23/17 | 10:29 | 26+ | 38.0550 | -122.4915 | San Pablo | Tidal Marsh | F |
| 4/4/17 | 09:10 | 27 | 38.0588 | -122.0381 | Suisun | Channel Bottom | - |
| 4/4/17 | 11:06 | 28 | 38.0748 | -122.0044 | Suisun | Tidal Marsh | - |
| 4/4/17 | 11:58 | 29+ | 38.1092 | -121.9922 | Suisun | Shoal | - |
| 4/18/17 | 15:13 | 35 | 38.0658 | -122.2065 | Suisun | Shoal | - |
| 5/25/17 | 11:53 | 39 | 38.0414 | -122.1091 | Suisun | Tidal Marsh | F |
| 5/25/17 | 12:57 | 40 | 38.0389 | -122.1126 | Suisun | Tidal Marsh | - |

*Date, time, and location (latitude and longitude) of fish and zooplankton samples (tow #) collected in northern San Francisco Estuary regions and habitats. + after a Tow # indicates zooplankton samples that were used as positive controls. Fish Present indicates whether longfin smelt or Pacific herring were found (F) in the fish sample or if neither fish was present (-).*

Two nets were towed to collect larval fishes and zooplankton concurrently: a 0.52 m diameter 505 μm mesh net with a filtering cod end for larval fish and a 0.35 m 150 μm mesh net with a filtering cod end for zooplankton. Each net was fitted with a General Oceanics flowmeter (Model 2030R, low-flow rotor). At each sample site two tows were conducted consecutively: one tow for molecular diet studies and a second for morphological diet studies. Larval fish and zooplankton from one tow were immediately preserved in 95% non-denatured ethyl alcohol (EtOH) and placed on ice for molecular diet analysis. The larval fish and zooplankton from the second tow were preserved in 2-4% formaldehyde (final conc., vol:vol) for use primarily in a separate study examining the distribution and abundance of larval longfin smelt and Pacific herring (Grimaldo et al., 2020), and for a study of the fish diets through morphological analysis. Upon return to the laboratory and within 12 hours, samples for molecular analysis were stored at −20 °C; EtOH was exchanged to fresh EtOH within 24 hours of collection to maintain sample integrity (Bucklin 2000).

### Fish Sample Processing

Fish in samples were sorted and identified to species. All longfin smelt were processed for sequencing. Up to 20 Pacific herring per sample were processed for analysis from each of four tows where both fish species were present and at least one species was relatively abundant (>10 individuals) (Table 2). We isolated selected fish from the sample, photographed and measured the entire body, and visually determined if prey were visible without dissection. Dissection tools were UV sterilized between samples, and wiped clean between fish within the same sample. For each larva, the entire digestive tract from the base of the gills to the anus was carefully separated from the body tissue using a fine probe and forceps. The digestive tract was then placed in a sterile tube containing 95% EtOH for DNA extraction.

**Table 2.** Samples where fish were collected, along with associated metadata on fish abundance, number of samples sequenced, and fish size.

| Date | Tow | Region | Habitat | Longfin Smelt |  |  |  |  | Pacific Herring |  |  |  |  |
| --- | --- | --- | --- | --- | --- | --- | --- | --- | --- | --- | --- | --- | --- |
|  |  |  |  | Fish<br>sample<sup>-1</sup> | Fish m<sup>-3</sup> | Seq. | Length<br>± SD<br>(mm) | Length<br>Range<br>(mm) | Fish<br>sample<sup>-1</sup> | Fish<br>m<sup>-3</sup> | Seq. | Length<br>± SD<br>(mm) | Length<br>Range<br>(mm) |
| 3/7/17 | 6 | Suisun | Tidal Marsh | 1 | 0.0184 | 1 | 7 | 7 | 0 | - | - | - | - |
| 3/7/17 | 13 | San Pablo | Channel Bottom | 21 | 0.1489 | 21 | 8 ± 2 | 7 – 17 | 36 | 0.255 | 20 | 12 ± 3 | 10 – 19 |
| 3/9/17 | 14 | San Pablo | Channel Bottom | 4 | 0.0487 | 3 | 10 ± 5 | 6 – 15 | 23 | 0.28 | - | 10 ± 4 | 8 – 22 |
| 3/9/17 | 15 | San Pablo | Tidal Marsh | 42 | 0.2973 | 21* | 11 ± 3 | 8 – 20 | 20 | 0.142 | 20 | 16 ± 6 | 10 – 28 |
| 3/9/17 | 16 | San Pablo | Channel Surface | 8 | 0.0584 | 8 | 15 ± 4 | 8 – 20 | 86 | 0.628 | - | 16 ± 5 | 10 – 24 |
| 3/9/17 | 17 | San Pablo | Channel Surface | 0 | - | - | - | - | 18 | 0.145 | - | 18 ± 4 | 9 – 22 |
| 3/22/17 | 20 | Suisun | Shoal | 46 | 0.3872 | 23* | 8 ± 1 | 6 – 10 | 0 | - | - | - | - |
| 3/22/17 | 21 | Suisun | Channel Bottom | 3 | 0.0216 | 3 | 13 ± 4 | 9 – 17 | 4 | 0.029 | - | 14 ± 3 | 11 – 16 |
| 3/22/17 | 22 | Suisun | Channel Bottom | 3 | 0.0229 | 3 | 8 ± 1 | 8 – 9 | 0 | - | - | - | - |
| 3/23/17 | 23 | Suisun | Shoal | 1 | 0.0076 | 1 | 7 | 7 | 13 | 0.099 | 13 | 16 ± 4 | 11 – 25 |
| 3/23/17 | 24 | San Pablo | Shoal | 25 | 0.1181 | 25 | 14 ± 3 | 9 – 20 | 349 | 0.128 | 20 | 20 ± 7 | 11 – 31 |
| 3/23/17 | 25 | San Pablo | Tidal Marsh | 0 | - | - | - | - | 43 | 0.142 | - | 16 ± 5 | 10 – 30 |
| 3/23/17 | 26 | San Pablo | Tidal Marsh | 2 | 0.0147 | 2 | 13 ± 3 | 12 – 15 | 87 | 0.638 | - | 18 ± 6 | 10 – 30 |

The total number (Fish sample<sup>-1</sup>) and abundance (Fish m<sup>-3</sup>) of fish collected in each sample, with the number of fish guts sequenced (Seq.), mean and SD of length (mm) and length range of fishes of each type. \* sequenced samples contained pooled extract from two fish guts

Fishes from four tows were analyzed for both dDNA and morphological diet analysis of longfin smelt larvae to compare results of each method. For morphological diet analysis, each formaldehyde-preserved larva was examined under a dissecting microscope and prey items in the entire digestive tract were identified, counted, and measured. Calanoid copepods, *Limnoithona tetraspina*, and adult *Acanthocyclops* spp. were identified to species. Other cyclopoids and harpacticoids were identified to order. Copepod nauplii were not further identified (labeled here as unidentified copepoda). Non-copepod prey were identified to the lowest taxonomic level possible.

### DNA Extraction

We desiccated the fish guts using a vacuum centrifuge at room temperature for a maximum of 1.25 hours. The desiccated guts were then resuspended in warmed buffer ATL (Qiagen) and vortexed, and guts from fish larger than 10 mm were pestle ground to facilitate tissue breakdown. One negative gut control was created along with each batch of gut extractions (n=3), consisting of a sterile microcentrifuge tube with the same clean ethanol that was used for the gut preservation, and processed in the same way as the guts. A standard protocol (*DNeasy Blood and Tissue kit, Purification of Total DNA from Animal Tissues*, Qiagen) was followed with an overnight incubation and addition of the recommended RNAse A step. Extracts were eluted twice with 50 μL (100 μL total) into a single tube and placed in a −80 °C freezer.

We then extracted DNA from a quantitative subsample from each zooplankton sample for comparison to the prey found in the diet. Each subsample volume was calculated so that 1) material remained in the sample for morphological identification and barcoding if needed later; 2) the volume of equivalent estuarine water extracted was equal across samples; and 3) the volume extracted was large enough to adequately represent the ambient zooplankton assemblage at densities typical of the northern estuary. Zooplankton subsamples were vacuum-concentrated onto bleach-sterilized 100 μm nitex filters to remove sample ethanol. Filters were transferred into 15 mL centrifuge tubes for extraction following the OMEGA EZNA soil DNA kit following the protocol for 250-1000 mg samples. This kit was chosen to minimize inhibition due to high concentrations of sediment in some samples. When the subsample was thick with material, it was split onto multiple filters for DNA extraction and the elution products from all splits of the same subsample were combined. In each of the four batches of DNA extractions a negative extraction control was included, consisting of a bleach-sterilized filter processed in the same way as the zooplankton samples.

After extracting quantitative subsamples from the zooplankton tow samples, we used a subset of zooplankton tows to generate positive controls. These came from zooplankton tows 12, 13, 15, 16, 20, 23, 26 and 29, and represented channel, shoal, and marsh habitats in both San Pablo Bay and Suisun Bay (Table 1). Each sample was sorted to isolate at least one individual of each unique organism found; organisms were then identified to the lowest taxonomic level, and transferred into a single vial containing all sorted organisms for the sample. DNA extraction of positive controls was performed using the same DNeasy Tissue Extraction kit (Qiagen) protocol as described above, including grinding the organisms in each sample with a sterile pestle prior to extraction.

In two fish samples, each containing > 40 longfin larvae (tows 15 and 20), we pooled DNA extracts from pairs of larvae from the same sample. A total of 111 extracts were generated for longfin smelt larvae. Each of the guts from the 73 herring was sequenced individually. Bulk zooplankton was sequenced from 32 zooplankton samples, including the samples collected where longfin smelt and Pacific herring larvae were present (“fish” zooplankton) as well as samples where these two larval fishes were not present in concurrent fish samples (“no fish” zooplankton); analyzing both “fish” and “no fish” zooplankton samples allowed us to assess for possible differences in the zooplankton prey assemblages where larval fishes were present.

### Library Preparation and Sequencing

We sequenced zooplankton and fish gut samples on two separate MiSeq sequencing runs to recover more sequences per sample from the smaller number of zooplankton samples, since plankton diversity was expected to be much higher in zooplankton samples than in fish guts, and to maximize sequencing coverage in the large number of fish guts. For each sequencing run, DNA in extracts was first amplified (PCR1) with the following universal metazoan primers (Table 3) that included standard Nextera Transposase Adapters (Illumina) to target the mtCOI gene: mjHCO2198 (modified from Geller *et al*., 2013), and mlCOIintF (Leray *et al*., 2013). The mjHCO2198 primers were modified from Geller *et al*., (2013) because we found the inosine bases to be incompatible with high-fidelity DNA polymerase. In addition to using the universal mtCOI primers, fish-gut extracts were amplified with an annealing-inhibiting primer (blocking primer; 10-fold higher concentration), which were designed for this study to reduce amplification of DNA from the fish (e.g., Vestheim & Jarman 2008): Lfs_COIBlk_668R for longfin smelt and Clp_COIBlk_668R for Pacific herring. Blocking primers included a phosphorothioate bond at the 3’ end to prevent exonuclease degradation of the C3-spacer by the high-fidelity polymerase.

**Table 3.**
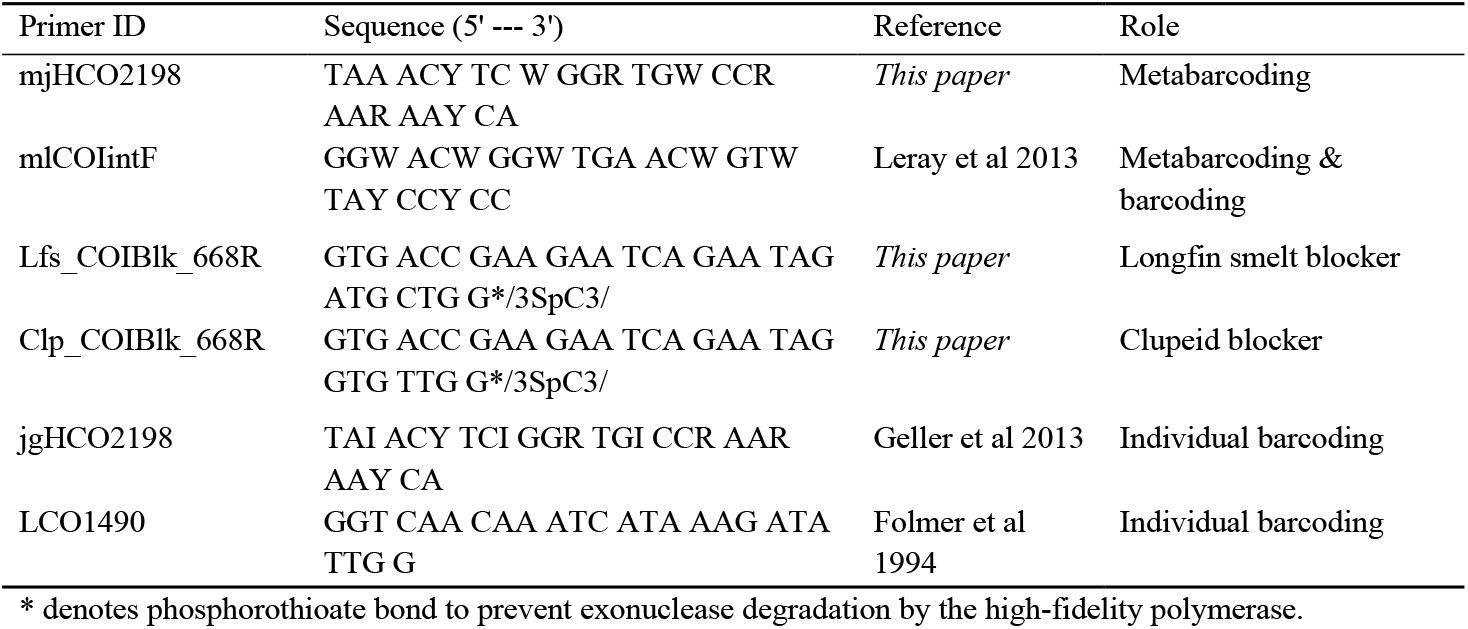
PCR and blocking primers used in this study.

The zooplankton assemblage and associated negative controls, including two PCR negative controls, were sequenced on the first sequencing run. Each reaction in PCR1 for the first sequencing run was prepared with triplicate PCR amplifications for each DNA extract with the following reaction setup: 2 μL of 5x Kapa fidelity buffer, 0.3 μL dNTPs (10 mM), 1 ng bovine serum albumin (BSA, to minimize inhibition), 0.3 μM of each universal primer, 0.2 μL Kapa HiFi Polymerase, and 1 μL of DNA extract, with nuclease-free H_2_O added to a total volume of 10 μL. The PCR 1 thermal-cycling protocol included initial denaturation at 95 °C for 3 minutes, followed by 25 cycles of 98 °C for 20 seconds, 46 °C for 30 seconds, and 72 °C for 15 seconds, with a final 72 °C extension for 4 minutes. Triplicate products from PCR 1 were pooled before indexing the products in the second PCR. The second PCR was performed with the following conditions: 5 μL of 5x Kapa fidelity buffer, 0.75 μL dNTPs (10 mM), 0.5 mM MgCl_2_, 1 ng BSA, 0.5 μM of each indexing primer, 0.5 μL of Kapa HiFi Polymerase, and 2.5 μL of pooled PCR1 product with nuclease-free H_2_O added to a total volume of 25 μL. The PCR2 thermal-cycling protocol included initial denaturation at 95 °C for 3 minutes, followed by 8 cycles of 98 °C for 30 seconds, 55 °C for 30 seconds, and 72 °C for 30 seconds, with a final 72 °C extension for 5 minutes. Gel electrophoresis indicated strong product amplification. We performed a single Serapure bead cleanup (Bronner *et al*., 2013; Rohland & Reich 2012) to remove remaining primers and dNTPs. After a Qubit HS assay was used to quantify all zooplankton products, equimolar concentrations of each barcoded sample (7 nM) were pooled to a single library tube. After denaturation, the final library was sequenced at the San Francisco State University Genetics Transcriptomics Analysis Core facility on an Illumina MiSeq platform using paired-end sequencing (MiSeq Reagent kit V2) with a 20% PhiX spike-in control to improve quality of low-diversity samples.

For the second sequencing run which included the fish gut samples, gut extraction controls, positive controls, and four PCR negative controls, we used Kapa HiFi Hotstart Readymix (Kapa Biosystems) to simplify the mastermix protocol, and amplified each DNA extract in triplicate in PCR1 with the thermal cycling conditions described above but with 35 cycles of the denaturation, annealing, extension steps, and the following reaction setup: 5 μL Kapa 2x HiFi Hotstart Readymix, 0.3 μM of each universal primer, 3 μM of the fish-specific blocking primer, 1 ng BSA, 2 μL DNA extract, and the remaining volume of nuclease-free H_2_O for a 10 μL reaction per replicate. Triplicate products from PCR1 were pooled, and PCR2 was performed to index each library with the following conditions: 12.5 μL of Kapa 2x HiFi Hotstart Readymix, 0.5 μM of each indexing primer, 1 ng BSA, and 2.5 μL of pooled PCR1 product, and an appropriate volume of nuclease-free H_2_O for a 25 μL reaction per sample. Bead cleanup was performed to remove both large non-target and small non-target DNA fragments (Bronner *et al*., 2013) from the amplified gut samples. The DNA quantity in indexed, bead-cleaned PCR products was normalized using the SequalPrep Normalization Plate Kit (Applied Biosystems) to obtain 250 ng PCR product per well. Five μL of each product were pooled to create the final library provided to UC Davis Sequencing facility. The final library was run on an Illumina MiSeq platform using paired-end sequencing (MiSeq Reagent kit V2) with a 35% PhiX spike-in control to improve the quality of the low sequence diversity expected in the sequenced fish gut samples.

### Sequence Analysis

Sequence data were processed with custom scripts written to analyze Illumina-generated metabarcoding data. Initial processing of these data employed primer trimming with cutadapt (v2.1), read pairing with pear (V0.9.11; Zhang *et al*., 2014), and following the DADA2 pipeline (v1.10.1; Callahan *et al*., 2016). The DADA2 pipeline filters out sequencing errors, dereplicates sequences, identifies chimeras, merges paired-end reads, and identifies amplicon sequence variants (ASVs, 100% identical sequence groups).

We performed further quality filtering of the complete set of ASVs in several steps. First, we looked at the distribution of sequence lengths in the entire set of ASVs. Sequences that were 313 bp in length were kept for further analysis to minimize false positives and remove spurious sequences from the dataset (size-based filtering). All remaining ASVs were annotated by BLASTn to the NCBI nucleotide database (downloaded 21 March 2019). The top hit (≥ 97.0% identity, ≥ 90% query coverage) was retained from this set, and any ASVs that remained with <97% identity were then processed with the MIDORI classifier (Machida *et al*., 2017) using the MIDORI-unique reference dataset (updated 21 February 2018) and RDP Classifier (Wang *et al*., 2007) with a cutoff of 80% bootstrap confidence. We assigned ASVs with < 80% bootstrap confidence to the “unknown” category. We then used taxonomy-based filtering on classified ASVs: any sequences classified as non-eukaryotes (98 ASVs, 952 reads) as well as non-actinopterygian chordates (3 ASVs, 37 reads) were excluded from the dataset (Suppl. Table 1), and sequences identified as the predator species (longfin smelt or Pacific herring) in the same species’ gut were also excluded from each corresponding gut sample (51,455 longfin smelt and 50,130 Pacific herring reads total). Remaining eukaryotic sequences not classified as metazoans were lumped into the “non-metazoan” group for further analysis.

The sequences were analyzed to identify gaps in the Genbank database for taxa known to be in the estuary and that may have been assigned to the “unknown” category. Cyclopoid and harpacticoid copepods were abundant in the plankton samples but poorly resolved in the database. We therefore sequenced DNA barcodes of individual copepods to fill these gaps. Individual cyclopoids (n=31) and harpacticoids (n=11) were selected from the bulk zooplankton samples and morphologically identified to the lowest possible taxonomic level. Individuals were cleaned of external debris, a voucher photo was taken of each organism, and total DNA was extracted following the standard Qiagen DNeasy DNA Extraction from Tissues protocol. DNA barcodes (mtCOI) were amplified in PCR with the following reaction setup: 10 μL MangoMix DNA Polymerase (Bioline), 0.25 μM of each primer, 5 μL DNA extract, and nuclease-free H_2_O for a 20 μL reaction. Two primer sets were used (Table 3). All extracts were first amplified with universal mtCOI primers LCO1490 (Folmer *et al*., 1994) and jgHCO2198 (Geller *et al*., 2013). Individuals that did not amplify with the first primer set were then tested with both metabarcoding primers used in this study. PCR products were cleaned (ExoSAP-IT, Affymetrix), ligated, cloned, and sequenced using standard protocols (ABI 3500 Perkin-Elmer capillary sequencer; BigDye v3.1). DNA sequences were compared to the NCBI Genbank database using BLASTn and any sequences that did not result in a >97% ID match were added to our local database and BLASTn-matched to ASVs in the current dataset.

### Statistical Analysis

Statistical analyses were performed in R (R Core Team 2019). Environmental variables were standardized and a Principal Component Analysis (PCA) was used to evaluate environmental variation among sampling events (*stats* R package). Differences in beta diversity across sample types were assessed with a non-metric multidimensional scaling (NMDS) analysis using Bray-Curtis dissimilarity on the ASV abundances, keeping taxa that were seen more than once in > 1% of samples (removed 5237 ASVs) to assess for similarity among the assemblages in the diets and zooplankton samples.

A Permutational Analysis of Variance (PERMANOVA) was used to assess differences in beta diversity of ASVs among fish species, tow number, sampling region (San Pablo Bay or Suisun Bay), and sampling habitat (channel surface, shoal, tidal marsh). A type III sum of squares PERMANOVA was performed on the normalized ASV abundances using the adonis function (vegan package; Oksanen *et al*., 2019) with 999 permutations, and tested for interactions among all terms. For this analysis, the ASV abundances were normalized by rarefying to the median sequencing depth of 228 found for fish gut samples. Tests for homogeneity of dispersion among groups, an assumption of PERMANOVA, were performed using betadisper on the Bray-Curtis distance matrix (phyloseq package; McMurdie & Holmes 2013) of sample ASVs for each group of interest: fish species, tow number, sampling region, and habitat.

We calculated the percentage frequency of occurrence (FO) of prey taxa in each sample type (longfin smelt, Pacific herring, and zooplankton). Percentage FO is based on presence or absence of a prey item and is the percentage of the total number of a sample type with a given prey item (Baker et al., 2014; Hynes 1950). The FO of longfin smelt prey identified through dDNA analysis were compared to the FO of prey identified through morphological diet analysis (full study to be published elsewhere) in the four fish samples specifically chosen for this comparison.

Relative read abundance (RRA) was calculated (as in Deagle *et al*., 2019) to assess the relative abundance of a species’ DNA in each sample. The RRA gives each gut or zooplankton sample equal weight in the view of overall sequence abundance across samples of each type. *RRA_i_* for food item *i* was calculated as:

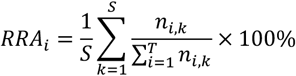

where *n_i,k_* is the number of sequences of food item *i* in sample *k*, *S* is the total number of samples, and *T* is the number of food items (taxa).

Differential abundance analysis (*DESeq2* and *phyloseq* R packages; Love et al., 2014; McMurdie & Holmes 2013) was used to assess differences in the relative sequence abundance of different taxa between zooplankton samples and diet samples. Differential abundance was calculated for the subset of taxa that overlapped between the fish diets and zooplankton assemblage. A likelihood ratio test (analogous to an ANOVA) was used to compare the relative abundance of each prey type (species-level or best identification) in the diet to that in the zooplankton assemblage with a maximum false discovery rate of 5%. The DESeq2 analysis accounts for differences in sequencing depth inherent in high-throughput sequencing, and when the observed sequence abundance is zero in one group, the likelihood ratio test utilizes a prior distribution on the log fold changes to provide differential abundance estimates (Love et al., 2014).

We used Schoener’s index (Schoener 1970; also known as Czekanowski’s index) to characterize overlap in diets between the two fish species (Feinsinger *et al*., 1981; Keppeler *et al*., 2015 Waraniak *et al*., 2019). Values of this index range from 0 (no overlap) to 1 (complete overlap) and values above 0.6 are considered biologically meaningful (Wallace 1981). Schoener’s index was calculated using the RRA values of each prey item for longfin smelt and Pacific herring.

## Results

Values of water turbidity, salinity, and chlorophyll were higher in some San Pablo Bay samples than in the Suisun Bay samples collected between February and April 2017. Across all sample regions and dates, water temperature ranged from 10 to 19 (median 14.4) °C, salinity ranged from 0.07 to 15.2 (median 0.2, Practical Salinity Scale), dissolved oxygen ranged from 8.3 to 13.9 (median 9.4) mg L^−1^, chlorophyll ranged from 0.5 to 9.9 (median 4.4) μg Chl L^−1^, and turbidity ranged from 9 to 153 (median 45) NTU. In general, water temperature increased in both regions over time and salinity increased over the first month of study in San Pablo Bay, from 0.1 on 28 February to 5.8 PPT on 23 March. In the samples where fishes were collected, water temperature ranged from 11 to 16 (median 14) °C, salinity ranged from 0.1 to 15.2 (median 1.2) PSU, dissolved oxygen ranged from 8.6 to 10.3 (median 9.5) mg L^−1^, chlorophyll ranged from 2.2 to 9.7 (median 7.7) μg Chl L^−1^, and turbidity ranged from 13 to 153 (median 85) NTU (Suppl. Figure 1).

The first two principal components of environmental data explained 75% of the variation among the samples (Figure 2). The first principal component (52% of variation) shows separation of samples primarily by dissolved oxygen (negative) and chlorophyll (positive). The second principal component (23% of variation) distinguishes sample salinity (negative) from oxygen, turbidity and chlorophyll (positive). Samples were grouped loosely by region (San Pablo Bay vs. Suisun Bay). Neither environmental variables nor sampling region were useful in distinguishing samples containing one or both species of fish from those without these fish.

**Figure 2:**
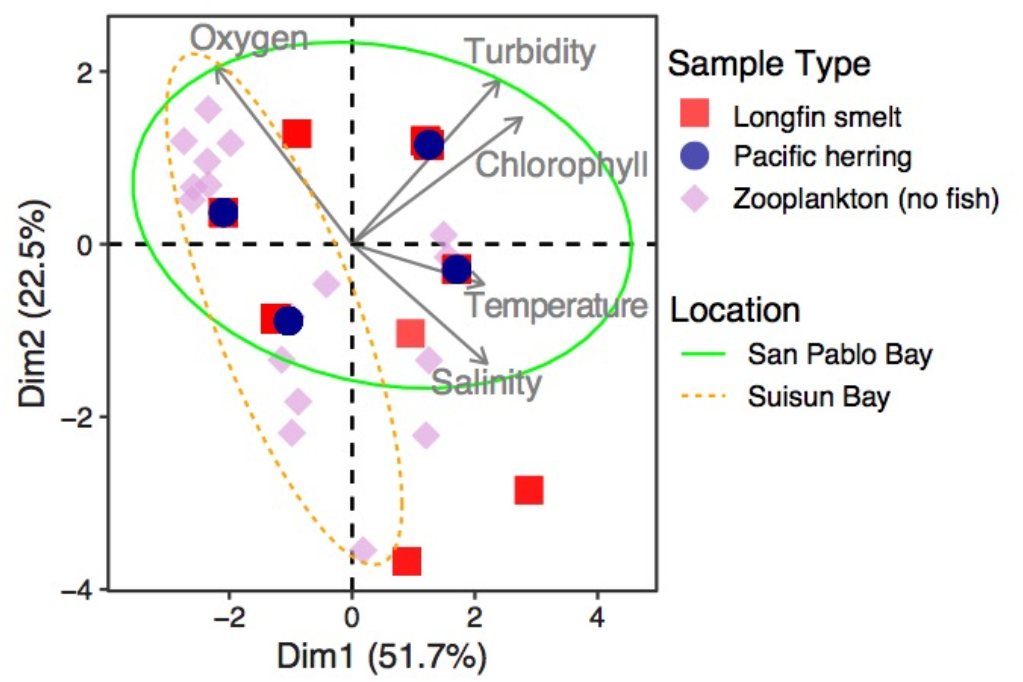
PCA biplot of environmental data showing samples containing longfin smelt (red square), Pacific herring (blue circle), or only zooplankton (pink diamond). Ovals indicate 95% confidence ellipses for sample groups in Suisun Bay (orange dotted line) and San Pablo Bay (light green line). Grey arrows indicate the environmental factors driving the axes of variation; salinity (PSU), water temperature (°C), chlorophyll-a (μg L^−1^), turbidity (NTU), and dissolved oxygen (mg L^−1^).

A total of 155 longfin smelt larvae were collected from 11 of the 32 sampling events between February and May 2017; only four samples contained > 10 longfin smelt (Table 2). To allow for a comparative analysis of diets, a total of 73 Pacific herring larvae were isolated from the four samples containing both species. Lengths of sequenced longfin smelt ranged from 6.2 to 20.0 mm (10.4 ± 3.6, mean ± SD), and lengths of sequenced Pacific herring were from 8.9 to 22.8 mm (16.1 ± 5.9, mean ± SD).

### Fish Diet and Community Metabarcoding

In total, high-throughput mtCOI amplicon sequence data were generated from 220 zooplankton and fish gut samples, six negative DNA extraction controls, and four negative PCR controls. A total of 10.6 × 10^6^ raw reads were recovered from the two sequencing runs. After DADA2 processing, size-based filtration, and taxonomic filtration, 4.2 × 10^6^ sequence reads remained containing a total of 9.6 × 10^3^ ASVs (Suppl. Table 2). Of the total sequence reads that remained, 82.7% were metazoan, 4.9% were non-metazoan, and 12.4% were classified as “unknown”. Of the total ASVs that remained, 41.3% were metazoan, 3.0% were non-metazoan, and 55.7% were classified as “unknown”.

Among the metazoan DNA sequences, predator DNA for each fish species accounted for 0.7% – 100% (median 73.5%) of the total sequence reads for each longfin smelt gut and 0.2% – 100% (median 82.3%) for each Pacific herring gut. There were 4 – 4,206 (non-predator) metazoan sequence reads (median 532) from each longfin smelt gut sample and 2 – 5,060 metazoan reads (median 824) from each Pacific herring gut sample. Sixteen (14% of the total) longfin smelt guts were considered empty (14 with 100% predator DNA, two with no DNA) and 10 (14% of the total) Pacific herring guts were considered empty (all with 100% predator DNA).

We recovered a 40-fold greater total reads in the zooplankton samples (4.0 × 10^6^ reads) than in the fish guts (0.1 × 10^6^ reads): sequencing depth was higher in the first sequencing run than in the second. Sequence diversity, as estimated by the number of ASVs, was also roughly 50-fold higher in zooplankton samples (9.47 × 10^3^ ASVs) than in fish guts (0.19 × 10^3^ ASVs). As a result, the average number of sequences per ASV was roughly equal between the fish guts and the zooplankton samples.

In positive control samples, DNA sequencing results provided equal or higher resolution of all morphologically identified taxa, with a few exceptions (Table 4). Several species were resolved with DNA that could not be identified to species using morphology: *A. americanus, A. robustus, A. vernalis, Acanthocyclops* sp., and another genetic group of unidentified cyclopoids. Multiple Diptera were resolved with DNA as well, including *Chironominae* sp., *Paratanytarsus grimmii*, and another genetically distinct but unidentified dipteran. The presence of *Oithona davisae*, ciliates, cumaceans, gastropods, nematodes, polychaetes, and rotifers, identified in the samples by morphology, could not be confirmed with DNA (Table 4). Many of these taxa are not well represented in the genetic database (NCBI) and thus may have been sequenced and lumped into broader groups such as the “unknown” group or unidentified Arthropoda.

**Table 4.**
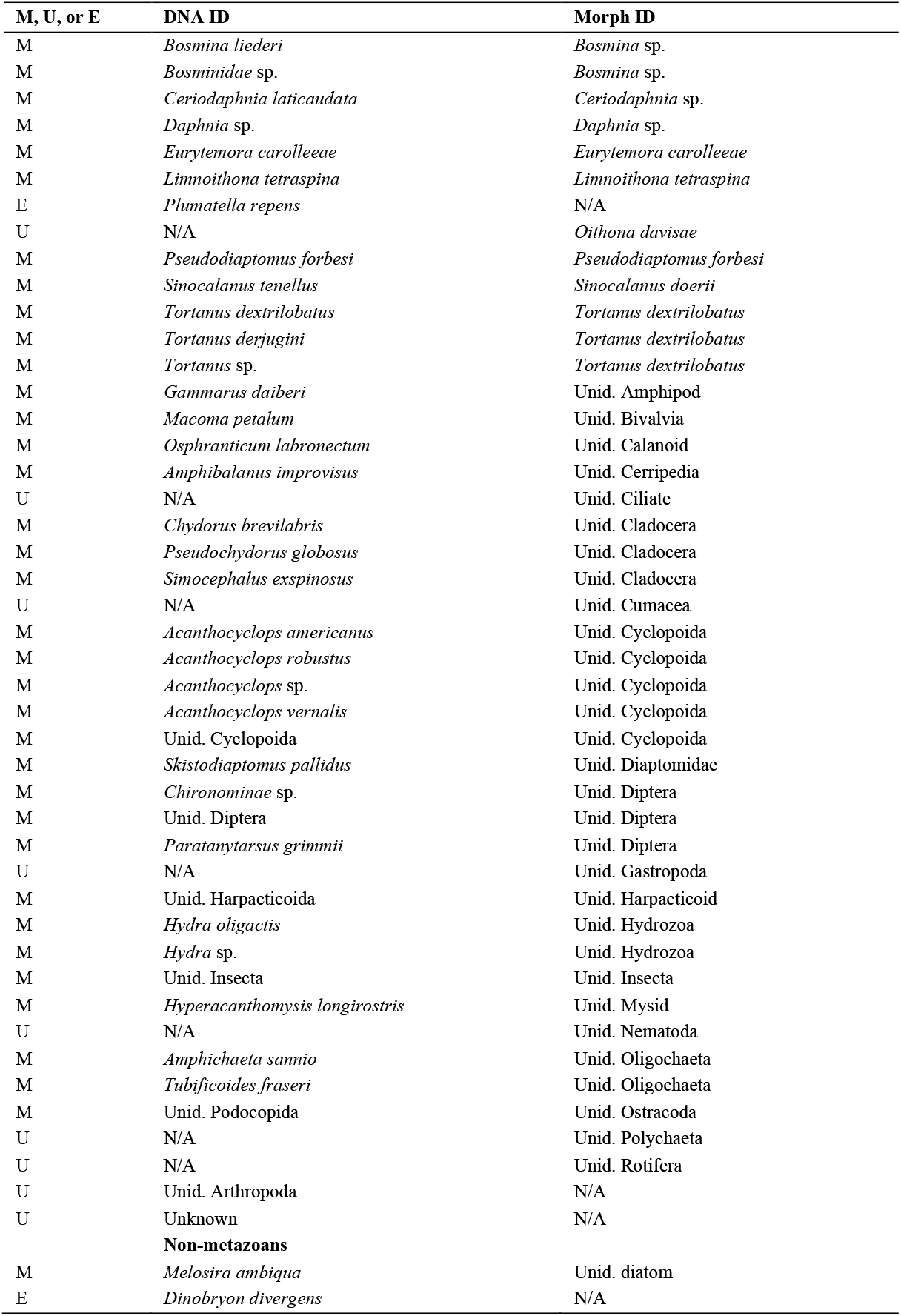
Positive controls comparing morphological ID (Morph ID) to the lowest taxonomic level with resulting DNA sequences (DNA ID). Match (M): DNA and Morph ID are a match, Unknown (U): ID does not have a clear match and corresponds to multiple types of organisms, Extra (E): not identified morphologically and DNA provided extra information. Unid: Unidentified to a higher level. N/A: ID for a column is unknown or not present.

| M, U, or E | DNA ID | Morph ID |
| --- | --- | --- |
| M | *Bosmina liederii* | *Bosmina* sp. |
| M | *Bosminidae* sp. | *Bosmina* sp. |
| M | *Ceriodaphnia laticaudata* | *Ceriodaphnia* sp. |
| M | *Daphnia* sp. | *Daphnia* sp. |
| M | *Eurytemora carolleeae* | *Eurytemora carolleeae* |
| M | *Limnoithona tetraspina* | *Limnoithona tetraspina* |
| E | *Plumatella repens* | N/A |
| U | N/A | *Oithona davisae* |
| M | *Pseudodiaptomus forbesi* | *Pseudodiaptomus forbesi* |
| M | *Sinocalanus tenellus* | *Sinocalanus doerrii* |
| M | *Tortanus dextrilobatus* | *Tortanus dextrilobatus* |
| M | *Tortanus derjugini* | *Tortanus dextrilobatus* |
| M | *Tortanus* sp. | *Tortanus dextrilobatus* |
| M | *Gammarus daiberi* | Unid. Amphipod |
| M | *Macoma petalum* | Unid. Bivalvia |
| M | *Osphranticum labronectum* | Unid. Calanoid |
| M | *Amphibalanus improvisus* | Unid. Cerripecta |
| U | N/A | Unid. Ciliate |
| M | *Chydorus brevilabris* | Unid. Cladocera |
| M | *Pseudochydorus globosus* | Unid. Cladocera |
| M | *Simocephalus exspinosus* | Unid. Cladocera |
| U | N/A | Unid. Cumacea |
| M | *Acanthocyclops americanus* | Unid. Cyclopoida |
| M | *Acanthocyclops robustus* | Unid. Cyclopoida |
| M | *Acanthocyclops* sp. | Unid. Cyclopoida |
| M | *Acanthocyclops vernalis* | Unid. Cyclopoida |
| M | Unid. Cyclopoida | Unid. Cyclopoida |
| M | *Skistodiaptomus pallidus* | Unid. Diaptomidae |
| M | *Chironominae* sp. | Unid. Diptera |
| M | Unid. Diptera | Unid. Diptera |
| M | *Paratanytarsus grimmii* | Unid. Diptera |
| U | N/A | Unid. Gastropoda |
| M | Unid. Harpacticoida | Unid. Harpacticoid |
| M | *Hydra oligactis* | Unid. Hydrozoa |
| M | *Hydra* sp. | Unid. Hydrozoa |
| M | Unid. Insecta | Unid. Insecta |
| M | *Hyperacanthomysis longirostris* | Unid. Mysid |
| U | N/A | Unid. Nematoda |
| M | *Amphichaeta sannio* | Unid. Oligochaeta |
| M | *Tubificoides fraseri* | Unid. Oligochaeta |
| M | Unid. Podocopa | Unid. Ostracoda |
| U | N/A | Unid. Polychaeta |
| U | N/A | Unid. Rotifera |
| U | Unid. Arthropoda | N/A |
| U | Unknown | N/A |
| **Non-metazoans** |  |  |
| M | *Melosira ambigua* | Unid. diatom |
| E | *Dinobryon divergens* | N/A |

We obtained sequences from 22 of the 31 cyclopoid and 11 harpacticoid copepods that were individually DNA barcoded, corresponding to 10 unique barcodes that we added to our database to improve resolution of “unknown” DNA sequences (Table 5). Ten individuals did not amplify, and 10 others resulted in sequences that did not correspond to the identified organism (Suppl. Table 3). Of the 10 additional unique DNA barcodes, one corresponded to the cyclopoid copepod *Limnoithona tetraspina* which was added to the database, two barcodes confirmed the presence of the two cyclopids, *Acanthocyclops americanus* and *Mesocyclops pehpeiensis*, and the remaining seven barcodes gave “unknown” ASVs a higher level of taxonomic resolution (Cyclopoida A, B, or Harpacticoida A, B) whose tentative morphological identification will require additional verification of species before they can be added to the public database (NCBI). In our study samples, 396 ASVs corresponded to *L. tetraspina*, two ASVs corresponded to Cyclopoida A, one ASV corresponded to Cyclopoida B, 137 ASVs corresponded to Harpacticoida A, and two ASVs corresponded to Harpacticoida B. The individual barcodes assigned to *A. americanus* and *M. pehpeiensis* corresponded to 379 and 1 ASV, respectively, for these two species in our study samples.

**Table 5.** DNA barcoded cyclopoid and harpacticoid copepods, with their tentative morphological identification (ID), the name applied to the ASVs that matched the morphological ID in this study, the best ID in NCBI, % identity of best NCBI match, NCBI best match accession number if > 97% identity, number of ASVs that matched that sequence type, and the number of individuals that were of that sequence type (n).

| Order | Tentative Morphological ID | Name Applied to ASVs | Best Database ID | % Match Best | Accession # | # >97% | n |
| --- | --- | --- | --- | --- | --- | --- | --- |
| Cyclopoida | *Limnoithona tetraspina* | *Limnoithona tetraspina* | Diptera | 81.4 | MF172405 | 396 | 5 |
|  | *Acanthocyclops americanus* | *Acanthocyclops americanus* | *Acanthocyclops americanus* | 98.8 | MG230154 | 379 | 1 |
|  | *Mesocyclops pehpeiensis* | *Mesocyclops pehpeiensis* | *Mesocyclops pehpeiensis* | 100 | MK159096 | 1 | 1 |
|  | *Homocyclops sp.* | N/A | Cyclopoida sp. | 99.4* | MG449991 | 0 | 1 |
|  | *Macrocyclus distinctus* | N/A | Cyclopoida sp. | 99.8* | MG449984 | 0 | 1 |
|  | *Acanthocyclops robustus* | N/A | Cyclopoida sp. | 99.8* | MG449984 | 0 | 1 |
|  | *Acanthocyclops robustus* | N/A | Cyclopoida sp. | 100* | MG449984 | 0 | 1 |
|  | *Diacyclops thomasi* or *A. capillatus* | N/A | Diptera | 80.1 | MF476244 | 0 | 1 |
|  | *Euterpina acutifrons* | Harpacticoida A | Hexapod | 74.1 | KJ083558 | 137 | 9 |
|  | *Pseudobradya sp.* | Harpacticoida B | *Metacyclopsina harpacticoida* | 79.5 | MH976658 | 4 | 1 |

\* indicates a high-level match (> 99%) to an organism with a low level of taxonomic ID in the database (order-level).

### Dietary and Assemblage Diversity

NMDS analysis shows the ASV assemblages in the longfin smelt and Pacific herring dDNA largely overlapped, while the zooplankton assemblages were tightly clustered in the middle of the plot (Figure 3). The tight clustering of zooplankton samples in the center suggests there were similar levels of diversity across zooplankton samples from different regions and habitats, while the fish gut samples were more spread out, suggesting there were more differences in diversity among different fish gut samples, in part due to fewer prey items sequenced in each gut sample, and with no discernible pattern. There was no separation between zooplankton assemblages where fish were present and where fish were not present. Seven outliers had to be removed from the NMDS analysis: six Pacific herring and one longfin smelt gut. Each outlier contained < 5 ASVs, five of which had fewer than 90 total reads, while the remaining two, both herring, had 1514 and 429 reads total.

**Figure 3.**
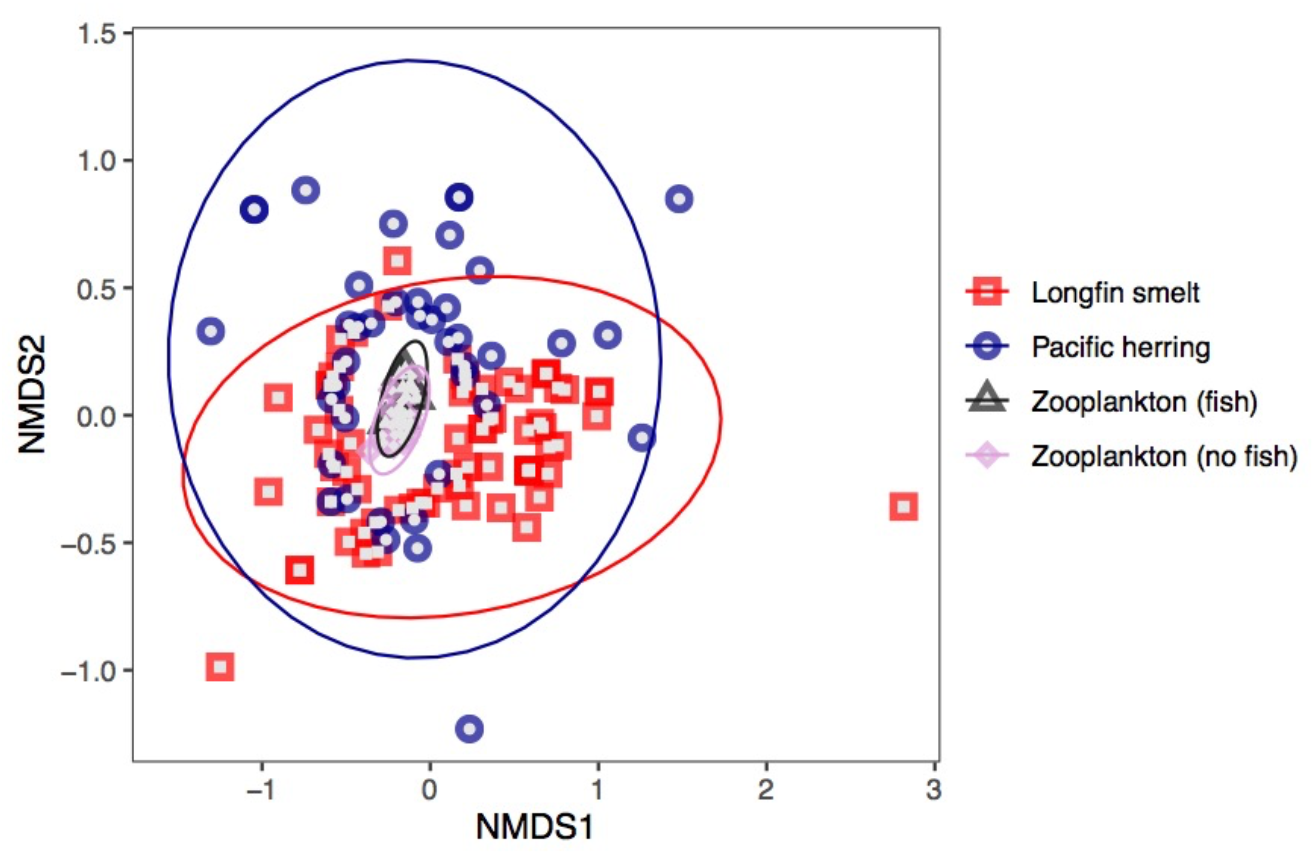
NMDS plot of normalized sequence diversity in each sample, with colors representing different types of samples: longfin smelt (individual or paired guts; red square), Pacific herring (individual guts; blue circle), zooplankton samples associated with fish (grey triangle), and zooplankton samples without fish (pink diamond). Ellipses represent 95% confidence groupings of each sample type.

Beta diversity tests (PERMANOVA) comparing the normalized ASVs in the dDNA of the two species suggest there may have been diet differences between the fishes (F_1,131_ = 2.66, p = 0.001) and between study regions (F_1,131_ = 3.69, p = 0.001), but there were also interactions between fish species and region (F_1,137_ = 3.61, p = 0.001). The significant interaction may indicate that there were both differences in diet between the two fish species as well as differences in diet of the same species in the two regions, but it is hard to tell with the statistical results alone. Differences among samples (tow number) and habitats were also tested, but the dispersion (variance) was not homogeneous when grouping prey assemblages by tow number or habitat (p = 0.001).

Schoener’s index of dietary overlap (alpha) calculated for longfin smelt and Pacific herring diets overall was 0.66, which is considered biologically meaningful (> 0.6) and supports the NMDS results showing largely overlapping sequence assemblages between the two species. For longfin smelt, we identified 116 ASVs of which 35 were classified with low confidence and categorized as “unknown,” representing 5% of the total longfin smelt diet sequences. For Pacific herring, we identified 99 ASVs of which 41 were classified as “unknown” and represented 11% of the total herring diet sequences.

Among ASVs classified to a known taxon, a variety of metazoan taxonomic groups were detected in the larval fish guts, including arthropods, chordates, cnidarians, echinoderms, and molluscs, and some non-metazoans. In total, 25 metazoan taxa were identified in longfin smelt guts, 16 in Pacific herring, and 195 taxa in both types of zooplankton samples (with and without fish) (Suppl. Table 4). Sequences classified as non-metazoans and “unknown” were found in all samples. We found 10 taxa in common among zooplankton collected with fish and guts of both fish species, 7 taxa unique to longfin smelt, 3 unique to Pacific herring, 13 unique to the zooplankton collected with fish, and 90 unique to the zooplankton collected with no fish. The 10 taxa found in all three fish-associated sample types included the copepods *E. carolleeae, A. robustus, A. americanus, Acartiella sinensis*, and *L. tetraspina*, the cladocerans *Daphnia* sp. and *Ceriodaphnia laticaudata*, the mysid *Neomysis* sp., an unidentified arthropod, and an unidentified cyclopoid (Cyclopoida C). Variation in prey taxa was high among individual fish, even among those collected in the same sample (Suppl. Figures 2 and 3).

The prey taxa found only in longfin smelt guts included several fish species, namely *C. pallasii* (Pacific herring), *Cottus asper* (prickly sculpin)*, Gasterosteus aculeatus* (threespine stickleback), and *Acanthogobius flavimanus* (yellowfin goby). Several prey taxa previously unreported in longfin smelt or herring diets included the copepods *Osphranticum labronectum* and *Mesocyclops pehpeiensis*, the insect *Liposcelis rufa*, the cnidarian *Hydra oligactis*, and the polychaete *Dasybranchus* sp. DNA identified as longfin smelt was found in herring guts; other unique prey included the barnacle *Balanus crenatus*, the bivalve *Potamocorbula amurensis*, the cladoceran *Ceriodaphnia* sp., the decapod *Palaemon modestus* and an unidentified insect.

### Relative Read Abundance (RRA)

Individual variation in RRA was high among sequences from individual fish and zooplankton samples (Suppl. Figures 2 and 3). When aggregated across all fish within each sample set (tow), sequences classified as “unknown” made up 6.2 – 58.7% of the RRA in longfin smelt diets, 6.7 – 29.3% of the RRA in Pacific herring diets, and 5.3 – 19.5% of the RRA in fish-associated zooplankton samples (Figure 4). A majority of metazoan sequence reads in guts of both fish species were from arthropods, including the copepods *E. carolleeae, A. americanus*, and *A. robustus*, with *L. tetraspina* important in Pacific herring.

**Figure 4:**
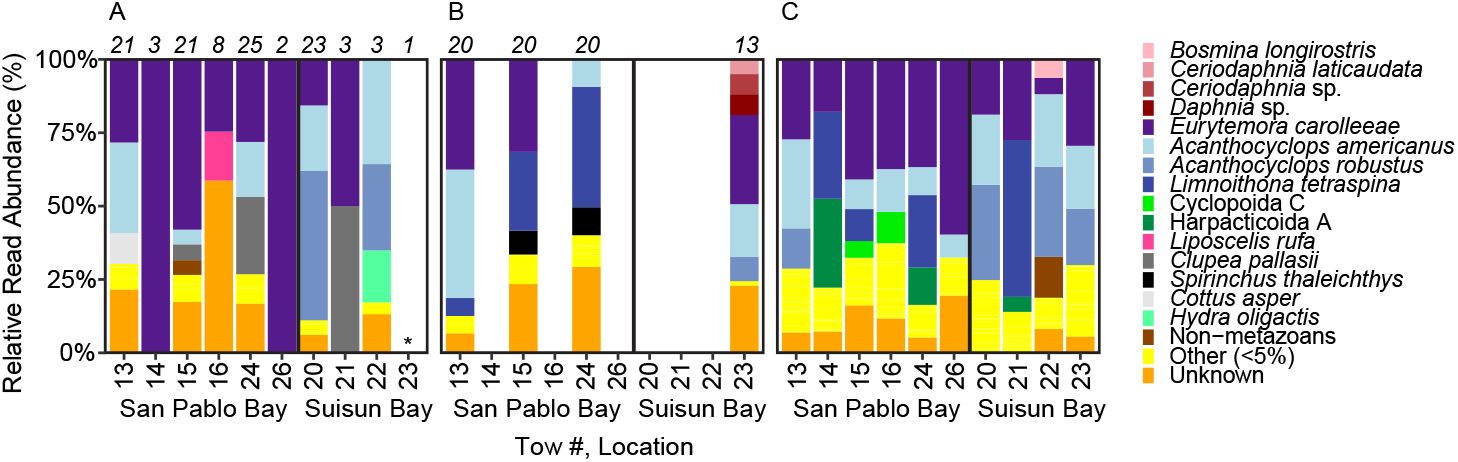
Relative read abundance (RRA %) in (A) longfin smelt (means for each tow), (B) Pacific herring (means for each tow), and (C) zooplankton samples associated with fish. Numbers above bars indicate the number of fish gut samples sequenced. “Other” includes prey IDs that contributed < 5% to the sample. “Unknowns” are those with < 80% RDP bootstrap confidence for ID. *empty gut.

The average RRA across all samples of each type shows which sequences were most abundant among all gut and zooplankton samples (Suppl. Table 4). In longfin smelt, *E. carolleeae* and *A. americanus* had the highest RRA values (33.0% and 17.5%, respectively) with *A. robustus* as third most abundant (14.7%). In Pacific herring, the three highest RRA values were from *E. carolleeae* (24.6%), *L. tetraspina* (20.3%), and *A. americanus* (18.3%). Sequences classified as “unknown” contributed an overall RRA of 17.0% in longfin smelt, 20.7% in herring, and 8.5% and 13.0% in fish-associated and no-fish zooplankton samples, respectively. Longfin smelt consumed relatively little *L. tetraspina* (1.9% RRA overall) even when they were abundant in the zooplankton sample, while the mean RRA for *L. tetraspina* in Pacific herring diets was 10-fold higher (20.3% RRA).

The highest RRA values in fish-associated zooplankton samples were also from the copepods *E. carolleeae* (30.1%), *A. americanus* (14.9%), *L. tetraspina* (13.1%), and *A. robustus* (11.4%). The highest RRA values in no-fish zooplankton samples were from the same taxa, only with lower relative abundances. An additional 166 less-common taxa were detected only in zooplankton samples with RRA values from < 0.001% to 5.3% in the fish-associated zooplankton samples and < 0.001% to 3.1% RRA in no-fish zooplankton samples (Suppl. Table 4).

Since copepods made up a majority of the known metazoan diet sequences, the RRA of copepods alone shows a higher-resolution view of the copepod prey of both larval fish species and highlights differences among samples (Figure 5). In general, presence of *E. carolleeae, A. robustus*, or *A. americanus* in the zooplankton sample usually coincided with presence and relatively high RRA in the diets of either fish. Other copepods in longfin diets at low RRA included *A. sinensis, O. labronectum, P. forbesi, L. tetraspina, M. pehpeiensis*, and Cyclopoida C (Suppl. Table 4).

**Figure 5:**
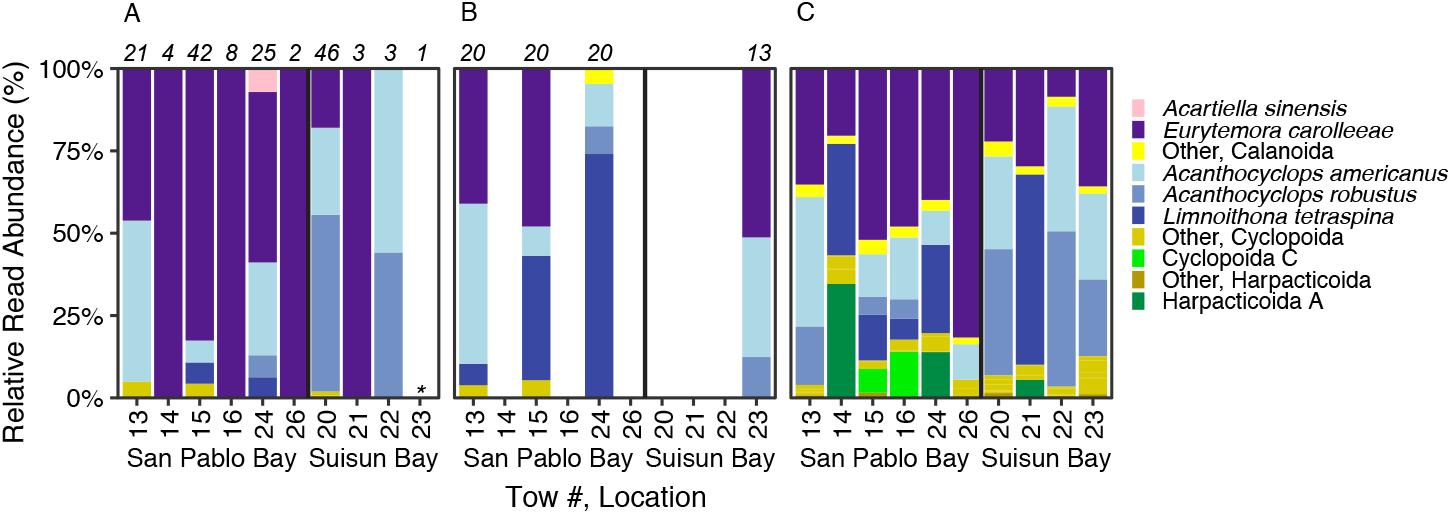
Relative read abundance (RRA %) of copepoda only, in (A) longfin smelt (means for each tow), (B) Pacific herring (means for each tow), and (C) zooplankton samples associated with fish. Numbers above bars indicate the number of fish gut samples sequenced. “Others” include prey IDs in each Order with < 5% contribution to the sample. *empty gut.

### Frequency of Occurrence (FO)

Copepods and fish were the most frequently detected prey in the longfin smelt larval diets (Figure 6). The prey with the highest frequency of occurrence in longfin smelt guts were identified as *E. carolleeae* (50.5%) followed by *A. americanus* (29.9%), *A. robustus* (19.6%), and *C. pallasii* (18.6%). *Eurytemora carolleeae* was also the most frequently detected prey in larval herring guts (44.4%) followed by *L. tetraspina* (30.2%), *A. americanus* (27.0%), and *S. thaleichthys* (12.7%) (Suppl. Table 4). “Unknown” DNA was present in 55.7% of longfin diets, 57.1% of herring diets, and all zooplankton samples.

**Figure 6:**
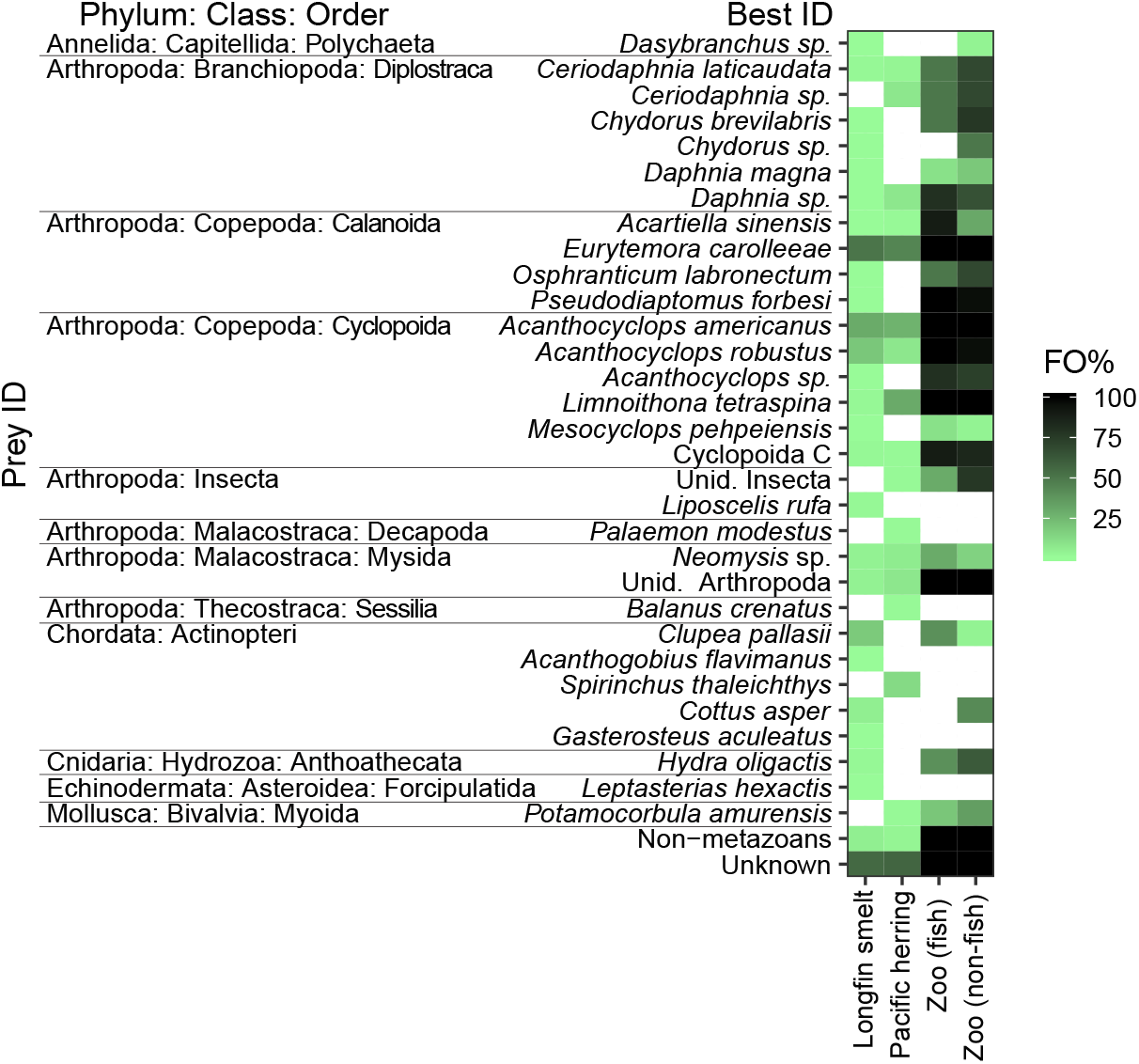
Heatmap of the frequency of occurrence (FO %) of taxa identified in the four sample types: longfin smelt, Pacific herring, zooplankton with fish (Zoo (fish)), and zooplankton without fish (Zoo (no fish)). Only prey taxa found in either longfin smelt or herring are shown here. White boxes indicate FO = 0%.

Several taxa were present in all fish-associated zooplankton samples (100% FO): *E. carolleeae, P. forbesi, A. americanus, A. robustus*, *L. tetraspina*, unidentified arthropods, and non-metazoans. Zooplankton samples not associated with fish samples had a similar representation of common taxa, except with a higher occurrence of unidentified insects with 76.9% FO, and lower FO for *P. forbesi, A. sinensis*, and *A. robustus*. The most common taxa present only in fish-associated zooplankton samples and not found in the fish guts included *A. vernalis* (100%), *Sinocalanus doerrii* (100%), Harpacticoida A (100%), Cyclopoida D (90%), and *Bosmina longirostris* (80%) (Suppl. Table 4). The most common taxa found in no-fish zooplankton samples and not found in fish diets included the Harpacticoida A (88.5%), *S. doerrii* (92.3%), *Bosmina liederi* (88.5%), *B. longirostris* (84.6%), and *Skistodiaptomus pallidus* (84.6%).

### Comparison of Morphological Diet to dDNA

Dietary DNA analysis of fish gut contents revealed a greater number of prey at a higher taxonomic resolution than morphological analysis in the subset of four longfin smelt diet samples used for this comparison (Figure 7). Taking into consideration differences in taxonomic resolution, all prey items identified by morphological analysis were also identified by molecular analysis with a few exceptions: rotifers, *S. doerrii*, and *T. dextrilobatus*, which were all present at low FO in the guts used for morphological analysis but not detected in the guts used for molecular analysis. For the two copepod species, there are few representatives for either species in the database (n=1 for *S. doerrii*, n=5 for *T. dextrilobatus*) so it is possible that the types present in our study are genetically distinct from what is in the database and could have been lumped into the “unknown” group as a result. Different species in the same genus were the best hit based on BLASTn found in other samples from our dataset: *S. tenellus* and *T. derjugini*. The result of these two being the top BLASTn hit likely resulted more from a lack of representatives of the two local species in the database, or from a few misidentified organisms in the database (see Discussion for more on this). The most frequently occurring species from morphological analysis, *E. carolleeae* (88.5% FO), was also the most frequently identified set of sequences from dDNA analysis (59.2% FO). Copepod groups reported morphologically as Unid. Copepoda, Unid. Calanoida, or Unid. Cyclopoida correspond to copepod nauplii, most of which would appear in the DNA database by species. As for the taxa that did not match well between methods, any rotifers in the DNA assemblage were likely classified as “unknown” because they are poorly represented in the database.

**Figure 7:**
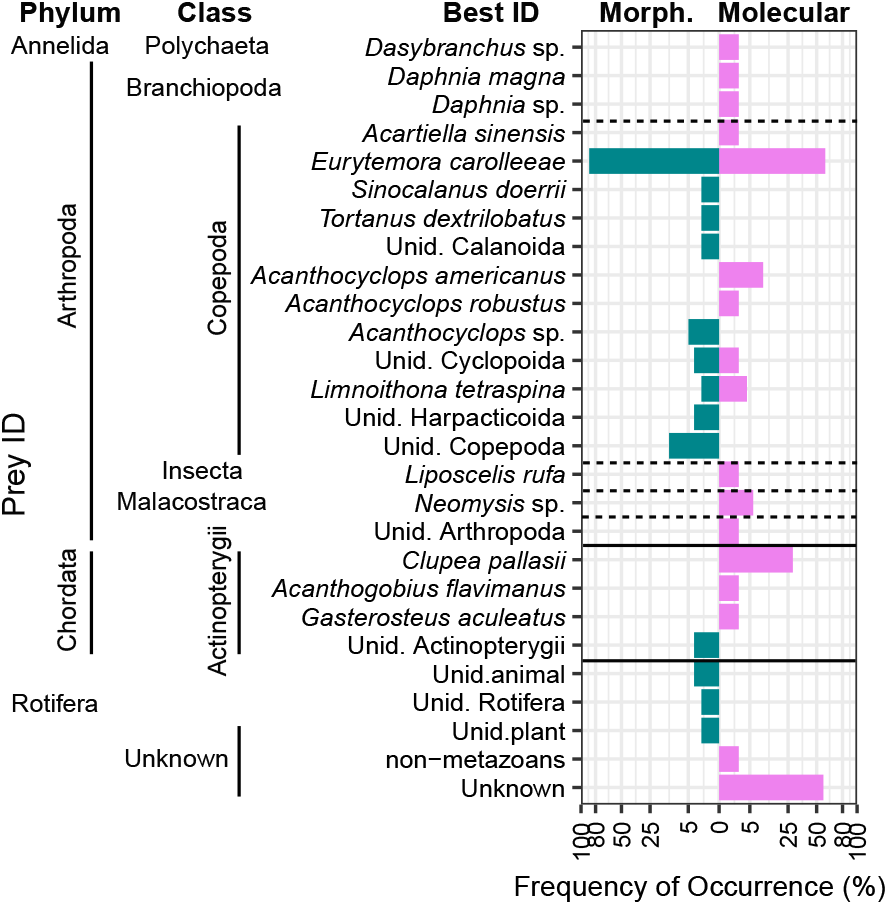
Comparison of the frequency of occurrence (FO %) of prey in longfin smelt diets obtained through morphological (morph.: turquoise bars) and molecular analysis (pink bars). Unid. indicates an unidentified organism of that general type. Note the x-axis is square-root transformed to expand the small values.

### Differential Abundance Analysis

Differential abundance (DA) analysis of longfin smelt dDNA found 24 of the 27 total prey types (including non-metazoans and unknowns) with DA values showing differences from zooplankton when using a false discovery rate of 5% (Figure 8A). The range of DA included *C. asper* at 22-fold higher in the diet than the zooplankton, to *L. tetraspina* at 0.001-fold of the value in the zooplankton. Of the prey types also found in zooplankton, *H. oligactis* was 2.9-fold higher in the longfin smelt diet, *M. pehpeiensis* was 4.4-fold higher, *D. magna* was 4.2-fold higher, non-metazoans were 0.04-fold lower, and “unknowns” were 0.02-fold lower. The highest values of DA were in prey types not found in zooplankton samples, including three of the four fishes, as well as *L. rufa, L. hexactis, Dasybranchus* sp., and *Chydorus* sp.

**Figure 8:**
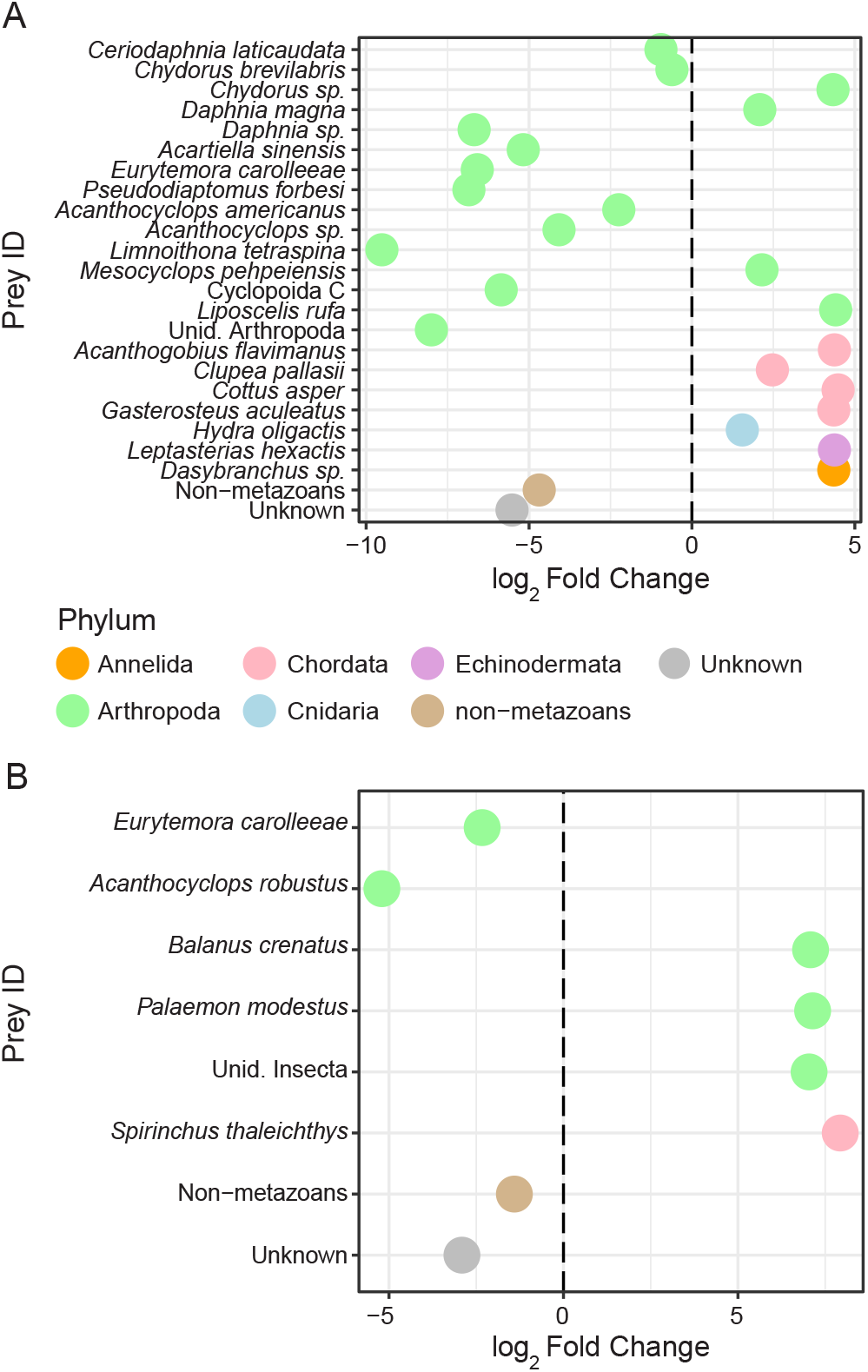
Differential abundance (log2 fold change) comparing the abundance of prey taxa from (A) longfin smelt guts and (B) Pacific herring guts with those from zooplankton samples. Prey types include only those present both in the guts and in the zooplankton. Positive fold change indicates prey relative abundance was higher in gut than in the zooplankton, negative fold change indicates prey relative abundance was lower in the gut than in the zooplankton.

The DA analysis for Pacific herring dDNA found eight of the 18 total taxa with DA values showing differences from zooplankton within the false discovery rate of 5% (Figure 8B). The range of DA included *S. thaleichthys* at 244-fold higher in the diet than in the zooplankton, to *A. robustus* at 0.03-fold the value in the zooplankton. Of the prey types also found in zooplankton, *E. carolleeae* was 0.2-fold lower in the Pacific herring diet, non-metazoans were 0.38-fold lower, and “unknowns” were 0.13-fold lower. The highest values were also in the prey types not found in the zooplankton samples, which include *S. thaleichthys, P. modestus, B. crenatus*, and Unid. Insecta.

## Discussion

This study is the first to apply dDNA analysis to elucidate the diets of both longfin smelt and Pacific herring larvae. Our results show that both calanoid (*E. carolleeae*) and cyclopoid (*A. americanus, A. robustus*, and *L. tetraspina* in herring) copepods were the most common and abundant prey for these two species of larval fishes, and diets of these two species largely overlapped. The most abundant prey sequences (as RRA) among zooplankton samples were also found in the fish guts, suggesting a general pattern of feeding on the most available prey, with some exceptions, such as *L. tetraspina*, which was infrequently consumed by longfin smelt despite its abundance. Twenty-five taxa were identified in the dDNA prey assemblage of the longfin smelt, and 16 in the dDNA prey assemblage of the herring, including all prey taxa known from prior morphological analysis in longfin smelt larvae and of prey taxa similar to those previously described in Pacific herring larvae (Bollens & Sanders 2004). While most prey taxa were arthropods and previously known prey, we also found many taxa that were not previously known to be consumed by these fish, some of which are soft-bodied species (e.g., the cnidarian *Hydra oligactis*, and the polychaete *Dasybranchus* sp.). We also provide DNA identification for species that are otherwise difficult to identify, including the copepod *M. pehpeiensis*, which was previously only identified to genus in the SFE. The zooplankton metabarcoding dataset generated here is likely to harbor additional discoveries of this nature that have not yet been identified, and will continue to provide information about the pelagic, benthic, and wetland-associated species present across the northern San Francisco Estuary.

### Common Prey

As expected, arthropods were the dominant prey type for the longfin smelt larvae, notably the copepods *E. carolleeae, A. americanus*, and *A. robustus*. Previous studies reported *Acanthocyclops* sp. as important prey (Hobbs et al., 2006) and at least three congeneric species are present in the estuary including *A. americanus, A. robustus*, and *A. vernalis*, which have been confirmed in the current study. Our study found that *A. americanus* and *A. robustus* were both present and common in the diet and in the zooplankton samples. *Acanthocyclops vernalis* was present in 100% of fish-associated zooplankton samples but uncommon (0.67% RRA; Suppl. Table 4) and was not consumed by either fish.

Arthropods were also the dominant prey type for Pacific herring larvae. Copepods were the most common and abundant prey sequences, with *E. carolleeae* and *A. americanus* among the most common copepod prey. Unlike longfin smelt, Pacific herring consumed moderate amounts of *L. tetraspina*. Given its greater abundance, *L. tetraspina* would be expected to play a role in the diets of fishes in the estuary. In fact, most other planktivorous fishes in the estuary do not commonly consume this species (Bryant and Arnold 2007; Slater and Baxter 2014), which is also supported by a prior study that found a low mortality rate in *L. tetraspina* (Kimmerer 2015).

### Uncommon Prey

Many less-common prey taxa occurred in the diets of both fish species that could be useful in identifying life history characteristics or feeding behaviors of the fishes *in situ*. For example, the presence of Pacific herring DNA in the guts of many longfin smelt larvae (19% FO) suggests that longfin smelt may have consumed herring eggs, which are adhesive and stick to substrates such as seagrass (Hay 1985). Pacific herring spawn every winter-spring in saline waters of the estuary (Watters *et al*., 2004); however, the extent of spawning further into the more brackish to fresh San Pablo and Suisun Bays has not been studied. The presence of herring DNA in longfin smelt diets from both Suisun and San Pablo Bays may indicate that longfin smelt larvae feed near substrates in these habitats, rather than in the open water. Moreover, a single herring egg (1.2 mm diameter, 125 μg C egg^−1^; Torniainen & Lehtiniemi 2008) contains about 38 times the organic carbon of a single adult *E. carolleeae* (3 μg C copepod^−1^; Pierson et al., 2016), making this a valuable supplement to the diets of the larval longfin smelt. Morphological diet analysis from the same field survey showed fish eggs in 6 of 551 longfin smelt gut samples, but eggs seen in morphological analysis were smaller than average-sized herring eggs and not identified to species (unpublished data) so may not have been herring eggs. Alternatively, herring DNA in the longfin smelt guts may have been from ingestion of herring feces or herring DNA present in detritus. Consumption of another fish species’ DNA through detritus or feces would be more likely if the species are schooling together, but there is currently no evidence supporting mixed schooling between larvae of these two species in the SFE, aside from being collected in the same larval fish samples. Schooling behavior in other fishes like northern anchovy *Engraulis mordax* and Atlantic silversides *Menidia* sp. is established between 10-15 mm (standard length; Hunter 1981; Hunter & Coyne 1982), so it is possible that the longfin smelt and Pacific herring were also forming schools during this study, given the dominant body sizes we sampled (Table 2). The detection of herring DNA in longfin smelt gut samples (or vice-versa) could have been from cross-contamination during library preparation or sequencing, which is a problem that can be hard to identify (e.g., Ballenghien *et al*., 2017). However, no Pacific herring-DNA sequences were found in our negative extraction or PCR controls, so it seems unlikely that this sort of cross-contamination was prevalent enough in our study to explain our results.

Some of the less common prey taxa hint at the possibility of longfin smelt larvae feeding on detritus. *Liposcelis rufa*, identified in diets in two longfin smelt, is in a group of terrestrial insects, psocids, commonly known as book or bark lice. Its presence in the diet of longfin smelt suggests that these fish could have eaten detritus from terrestrial runoff or wind-blown debris. In morphological analysis of longfin smelt larval diets, unidentified plant material occurred in a small number of smelt diets (Figure 7), also suggesting that smelt may have fed on detrital aggregates or near seagrasses. Detritus and associated organisms can be an important source of food for many estuarine and neritic organisms, from zooplankton to fishes (Adams 1976; Harfmann *et al*., 2019; Heinle *et al*., 1977). Future work amplifying the dDNA with additional primers that target different types of organisms such as plants and microbes could be useful in identifying habitats that support longfin smelt.

We found that 15 of the 25 (60%) identified taxa in longfin guts were not a part of the herring diet, and most of the unique prey were also relatively uncommon (Figure 6). One of the unique prey items in longfin smelt is also the first report of the copepod *M. pehpeiensis* in the SFE. Seven unique prey taxa in the longfin smelt were also not found in the fish-associated zooplankton and occurred at a low FO and RRA in four or fewer of the individual longfin-diet samples. These unique prey taxa included three fishes: *C. asper* (5.2% FO), *G. aculeatus* (1.0% FO) and *A. flavimanus* (1.0% FO). The presence of fish DNA in the longfin smelt larval diet most likely reflects occasional ingestion of the eggs of these species, or is evidence of the consumption of feces or detrital aggregates as discussed above for consumption of herring.

Six prey taxa in the Pacific herring diet were absent from longfin smelt. These included three which were likely ingested as meroplanktonic larvae: *P. amurensis, B. crenatus*, and *P. modestus* (Figure 6). *Potamocorbula amurensis* is an introduced clam that has decimated phytoplankton biomass throughout the SFE since its introduction in the 1980s (Alpine & Cloern 1992; Carlton *et al*., 1990; Kimmerer *et al*., 1994; Nichols *et al*., 1990). *Balanus crenatus* is a widespread acorn barnacle in the North Pacific and North Atlantic that has planktonic larvae (WoRMS 2020). *Palaemon modestus* (syn. *Exopalaemon modestus*) was introduced around 2000 (Brown & Hieb 2014) and has become a dominant component of the pelagic nekton assemblage in the freshwater areas of the Sacramento-San Joaquin Delta (Feyrer *et al*., 2017).

Prior studies of the diets of diverse fish larvae found differences in prey among families of fishes: some fed much more frequently on cyclopoid copepods, while others fed primarily on calanoids or specialized on other prey such as chaetognaths (Sampey et al.2007). The difference in assemblages of unique prey between the two fish species suggests that herring may forage in different sub-habitats or may be better equipped to detect and capture meroplankton larvae and cyclopoids than longfin smelt, but further work would be required to test this hypothesis.

### Non-metazoans and Unknowns

Non-metazoan taxa may be important prey for some fishes. In fact, the prior study of Pacific herring larval diets in the Estuary found a high proportion of tintinnid ciliates in fishes collected in the Central Bay (Bollens & Sanders 2004). Other studies have found *Tintinnopsis* spp. and *Eutinnus neriticus* to be the most abundant tintinnids throughout the SFE (Ambler et al., 1985) but these were not found in our fish diets or zooplankton samples. Six taxa identified to a high level of confidence in the diets of the two fishes were lumped into the non-metazoan category, five of which were found in the longfin smelt. These included a mixotrophic flagellate *Poterioochromonas malhamensis*, an amoeba that is a known parasite of fish *Cochliopodium minus* (Dyková *et al*., 1998), a lichen *Parmotrema stuppeum*, a centric diatom *Cyclostephanos* sp., and a fungal plant pathogen *Rhizoctonia solani*. The Pacific herring guts contained the mixotrophic flagellate *P. malhamensis* as well as a centric diatom *Skeletonema potamos*. The zooplankton samples included a range of more common taxa including some phytoplankton (*Melosira ambiqua, Thalassiosira pseudonana, Ditylum brightwellii*), a wheat-associated fungus (*Blumeria graminis*), a soil fungus (*Penicillium sclerotiorum*), and an aquatic fungus (*Tetracladium marchalianum*). *Melosira ambiqua* was the most common non-metazoan, found in all no-fish zooplankton samples and in 67% of fish-associated zooplankton samples. However, since the sequencing primers were chosen to target metazoan prey of the larval fishes, amplification of the phytoplankton, protist, amoeboid, or fungal groups is likely incomplete. The representation of many common non-metazoans in the DNA database is also currently poor and it is likely that some “unknown” DNA in the fish diets and environment corresponds to known species in the estuary, like the tinitinnid ciliates noted above.

Many sequences could not be identified to a reliable level from comparison to existing genetic databases (55.7% of ASVs, 12.4% of sequences), and further work is needed to determine what those prey are and whether they are important or informative prey items. A substantial number of the ASVs are likely to correspond to organisms that have not yet been identified and barcoded. For example, public mtCOI DNA sequences exist for only 61.2% of North American aquatic invertebrate genera (Curry *et al*., 2018).

### False Negatives

In addition to the above-stated uncertainties in identification due to “unknown” DNA in the samples, there is also a chance for false negatives. In this study, whole guts were used for DNA extraction and blocking primers were used to reduce the amplification of predator DNA. In initial tests validating the efficacy of the blocking primers developed here, blocking primers reduced amplification of predator DNA in a longfin smelt sample from 85% of the total reads to 18% of the total reads and resulted in the amplification of more prey types in the sample with blocking primers, though at a low number of reads. A similar result occurred with the use of clupeid blocking primers; we observed both a reduction in amplification of herring DNA (from 72% herring DNA to 18% herring DNA) as well as amplification of a broader range of less abundant prey from the gut.

Despite the use of primers meant for amplification of diverse metazoan taxa in our study, sometimes called “universal” primers, mtCOI genes of some taxa are not amplified effectively. For example, the mtCOI DNA barcode is notoriously difficult to amplify in cyclopoid copepods (Cepeda *et al*., 2012) as well as in nematode worms (Creer *et al*., 2010), and some neogastropoda and cardiida (Zhang *et al*., 2018). By individually DNA barcoding morphologically identified copepods, we found that even the “universal” metabarcoding primers used here did not successfully amplify some of the cyclopoid taxa in our samples, including individuals tentatively identified as *Acanthocyclops capillatus, Cyclops scutifer*, and *Eucyclops elegans* (Suppl. Table 3). Therefore, the full diversity of cyclopoid copepods and other taxa noted above is likely underestimated by this study. Taxonomic gaps due to primer bias could be filled by sequencing additional marker genes (e.g., Clarke *et al*., 2017; Zhang *et al*., 2018).

It is possible that the mtCOI gene primers applied here combined with strict removal of amplicons outside of the target 313 bp may have removed additional novel items in the larval fish diets. There were 7597 ASVs and 4.3 × 10^5^ sequences that passed the stringent DADA2 quality checking but did not fit the target amplicon length (313 bp; Suppl. Figure 4). A majority of these (99% in both species) would have been classified as “unknown” in our study with the classification methods applied here: Some of these had low-level matches to mtCOI sequences (e.g., < 97% identity to a sequence in the database), while others had no matches at all to sequences in the database. Within the remaining sequences that did not equal the target length of 313 bp and had a high-level (≥ 97% ID) match to a database sequence, a majority of these did not match the mtCOI gene. Within this subset of sequences that matched the mtCOI gene that were longer or shorter than 313 bp we found sequences that were likely rotifers and diatoms in the zooplankton samples, and land plants and a flatworm in the diet samples. Unfortunately, in order to exclude the majority of non-target gene sequences, we knowingly excluded a relatively small number of possibly real mtCOI sequences. Amplification of non-target genes is not surprising: Collins *et al*., (2019) reported extensive mis-priming of multiple primer pairs developed for metabarcoding mtCOI. Despite the known limitations of primer binding sites for metabarcoding the mtCOI gene, the benefits of species-level identification and the growing database of barcoded organisms means this gene remains the best choice for community metabarcoding studies on metazoans (Andujar *et al*., 2018).

Fourteen percent of stomachs were empty in each of the two fish species we analyzed. This is similar to rates seen through morphological analysis of diets in juvenile longfin smelt which found 13 – 21% of stomachs to be empty in Suisun Marsh (Feyrer *et al*., 2003). In a dDNA study on diets of post-larval clupeids in Tosa Bay, Japan, the researchers observed no empty guts using primer sets targeting genes with lower taxonomic resolution (Hirai *et al*., 2017), but morphological analysis of clupeid larval diets can result in up to 70% of fishes with empty guts. (Morote *et al*., 2010). Given the strict sequence classification and size-based exclusion used in our study as noted above, it is likely that some of our empty stomachs contained low levels of unique prey: re-sequencing the fish guts targeting an additional gene (e.g. 18S rRNA) may help reveal a broader range of taxonomic groups in the fish diets.

### A Case of Mistaken Zooplankton Identities

One of the more challenging aspects of eukaryotic metabarcoding is obtaining reliable identification of DNA sequences; this study is no exception. While we have made efforts to verify the DNA identity of the prey taxa described in this paper, some uncertainties remain. This remains to be solved, in part, because it can be hard to determine if the organism in the DNA database was properly identified before sequencing and thus, we end up having to make a judgement on the “real” DNA-based identity of an organism. For example, prior studies identified and described *Tortanus dextrilobatus* and *Sinocalanus doerrii* shortly after they were introduced to the upper Estuary (Orsi et al.1983; Orsi and Ohtsuka 1999). However, the best matches of our DNA sequences to the database for these genera were *T. derjugini* and *S. tenellus*, respectively for some ASVs, while *T. dextrilobatus* was the closest DNA match for other ASVs. Upon further inspection, alignment of DNA sequences from the Genbank database for all sequences corresponding to these four taxa suggests that a few misidentified organisms exist in the database, and that the original identifications (*T. dextrilobatus* and *S. doerrii*) are correct.

Another currently unsolved mystery lies in our *Neomysis* sp. sequence. In addition to other mysid genera, there are two *Neomysis* species described in the SFE: *Neomysis japonica* and *N. kadiakensis*. We recovered six ASVs that were assigned with high confidence to *Neomysis japonica* in our initial analysis. Upon further inspection the highest BLASTn hit for all six ASVs was to *N. japonica* but only at 93.5% ID (Suppl. Table 5). At that level, the sequence was assigned using RDP classifier, but with that level of match with BLASTn it is likely that it is a mysid, but probably not *N. japonica*. For now, we assign it as *Neomysis* sp. but acknowledge that it could be one of the other mysids of the SFE not presently in the NCBI database (see Suppl. Table 5 for other special cases).

### Conclusions

One aim of this study was to assess whether previously presumed food resources in the larval fish diets match what can be seen with a higher-resolution view of the prey field using dDNA, so that we can better assess the extent to which declining food resources are responsible for declines in fish abundance in the SFE. Overall, we found the prey assemblages in longfin smelt larvae and Pacific herring were similar and broadly reflect previous knowledge and concurrent morphological analysis of important prey (calanoid and cyclopoid copepods, primarily *E. carolleeae* at salinities in this study). In our study, both species relied on *E. carolleeae, A. americanus*, and *A. robustus* as dominant prey taxa in multiple samples. A key result of our study was that herring consumed *L. tetraspina* and other prey taxa that were not common prey for longfin smelt. These other prey taxa may provide an important source of food when larger calanoid copepods are not abundant, although other planktivorous fishes in the estuary also consume few *L. tetraspina* (Bouley & Kimmerer, 2006; Bryant & Arnold, 2007; Slater & Baxter, 2014; Sullivan *et al*., 2016) despite its numerical dominance in low-salinity regions (Bollens *et al*., 2011).

We examined larvae of these two species primarily in shallow, nearshore habitats and wetland channels, in addition to the fish collected in deeper channels. Previous studies show shallow habitats support larval longfin smelt (Grimaldo *et al*., 2017; 2020; Lewis *et al*., 2019). Current and planned wetland restoration (California Wetlands Monitoring Workgroup 2020) may provide more foraging opportunities for both fish species in these critical habitats, through greater contact with protective shelter and greater zooplankton biomass in habitats not readily colonized by clams. The declines in fish abundance in the estuary as a whole (Sommer *et al*., 2007) likely reflect the more or less parallel declines in the abundance of appropriate food for their larvae, notably the larger calanoid copepod *E. carolleeae*, which declined after the introduction of *P. amurensis* (Kimmerer *et al*., 1994).

In general, there were a greater number of species (ASV) sequenced from zooplankton samples than the fish diets. There was high variability in the prey taxa in individual fish guts (Suppl. Figure 2) and how much prey DNA was present. This may be a result of some of the fishes exploiting areas of patchy food availability and thus having different prey species in their diets. Larval fishes that exploit patches of concentrated food, such as in surface slicks in the open ocean, have higher survival likelihood and better body conditions than fishes found outside of concentrated food patches (Gove et al., 2019; Hunter 1981). Fish larvae are also visual feeders, and feeding is usually confined to the daylight hours due to limited visual abilities during the first few weeks of life; a prey item therefore has to be relatively nearby to be detected, and does not have to be motile (Hunter 1981). Feeding in a turbid estuary may present larval fishes with additional challenges but this turbid environment may also may provide the larval fishes with protection from visual predators (Boehlert & Morgan 1985; Sirois & Dodson 2000).

Larval fish growth rates depend strongly on food supply (Pepin *et al*., 2014). laboratory growth rates of Larval plaice and Atlantic herring reached maxima at food densities (copepod and *Artemia* nauplii) of ~ 500 prey L^−1^ in larval plaice and Atlantic herring in the laboratory (Kiorboe & Munk 1986; Wyatt 1972). Other clupeid species’ larvae also survive better at prey densities of up to 1000 – 4000 copepods L^−1^ in the laboratory (reviewed in Hunter 1981). In monitoring data taken in our sampling regions during springs of recent wet years (2006, 2011, this study 2017; “20-mm” survey, Dege and Brown 2004), calanoid copepod abundance (mostly juveniles and adults; nauplii undersampled) had a median of ~ 6 and rarely reached 10 copepods L^−1^, while the cyclopoids (mostly juveniles and adults; nauplii undersampled) reached about 1 copepod L^−1^. Thus, prey abundances in the northern SFE are one to two orders of magnitude lower than what seems to be required for high growth rates and high survival of most fish larvae. Our study also found that not all copepod prey are considered equally important to these fishes, as longfin smelt do not consume the abundant *L. tetraspina*, while Pacific herring seem to feed more generally. These results paint a disheartening picture of the foodweb support available to larval fishes in this region of the SFE and echo the need for ongoing restoration efforts to help enhance foodweb resources and protective habitat for fishes in the estuary.

Our findings illustrate the power of DNA-based methods to reveal feeding patterns and novel zooplankton diversity undetectable through complementary morphological analysis. We revealed larval fish feeding on two of the three *Acanthocyclops* spp., which are otherwise difficult to identify, found several fish species’ DNA in the larval fish diets, and identified larval meroplankton to the species level in the diets of the herring. These are just a few examples of the power of DNA-sequencing methods in studies of aquatic food web interactions. We also found at least one aquatic species not previously described in the SFE, the copepod *M. pehpeiensis*. This work provides a baseline for the genetic diversity of zooplankton in regions of San Pablo Bay and Suisun Bay in the northern SFE as of 2017, and a snapshot of the diets of two larval fishes during a wet year in 2017. Future research would benefit from targeted DNA barcoding of additional potential prey taxa that has poor representation in the NCBI database, such as meroplankton larvae and local aquatic-associated insects, and from deeper studies into the DNA recovered from unidentified arthropods, insects, and “unknown” DNA sequences.

## Acknowledgements

Funding was provided by the Sea Grant Delta Science Fellowship program and the State and Federal Contractors Water Association grant #18-02, an NSF-REU fellowship for A. Katla at San Francisco State University, grant P1696013 from the California Department of Fish and Wildlife to San Francisco State University. Larval fish trawls were collected under California Fish and Wildlife Prop 1 Grant study 2081; CESA Scientific Collecting Permit #4086 to ICF Inc. Funding for the San Francisco State University MiSeq sequencer was provided by NSF Award #1427772. Fish diet sequencing was carried out at the DNA Technologies and Expression Analysis Cores at the UC Davis Genome Center, supported by NIH Shared Instrumentation Grant 1S10OD010786-01. We thank the UC Davis Fish Conservation and Culturing Laboratory facility for providing longfin smelt adults to aid in primer design, C. Brennan for expertise in fish taxonomy, M. Conrad for assistance with library preparation, and F. Feyrer for providing input along the development of the project.

## Supplementary Tables

**Supplementary Table 1.** Non-actinopterygian chordates identified with BLASTn, with >= 97% identity to Phylum Chordata.

| Sample Type | SampleID | ASV | % ID | E-value | Phylum | Class | Order | Family | Genus | Species | Accession | Reads |
| --- | --- | --- | --- | --- | --- | --- | --- | --- | --- | --- | --- | --- |
| longfin smelt<br>mixed | LLF58 | ASV_8302 | 99.7 | 1.18E-155 | Chordata | Mammalia | Carnivora | Felidae | Felis | Felis silvestris lybica | KP202275.1 | 24 |
| zooplankton | Z27 | ASV_3600 | 100 | 2.77E-157 | Chordata | Mammalia | Primates | Hominidae | Homo | Homo sapiens | MK618711.1 | 8 |
| longfin smelt | LLF35 | ASV_22093 | 100 | 2.77E-157 | Chordata | Mammalia | - | Suidae | Sus | Sus scrofa | MF398983.1 | 5 |

**Supplementary Table 2.** Summary of Reads and ASVs in different sample types (longfin smelt guts, Pacific herring guts, zooplankton and controls), at each step of quality checking and data filtration (DADA2, size-based filtration, and taxonomy-based filtration, performed sequentially), and in different levels (metazoans, non-metazoans, unknowns) after taxonomy-based filtration.

|  | Longfin<br>Smelt Guts | Pacific<br>Herring<br>Guts | Zooplankton | Controls | Total |
| --- | --- | --- | --- | --- | --- |
| **# Samples Sequenced** | 111 | 73 | 36 | 10 |  |
| **Raw Reads** | 1,284,353 | 1,251,539 | 7,972,137 | 97,620 | 10,576,006 |
| **DADA-filtered Reads** | 222,970 | 191,049 | 4,559,113 | 3,835 | 4,976,967 |
| *DADA-ASVs* | 1,716 | 2,945 | 28,457 | 65 | 32,736 |
| **Size-filtered Reads** | 130,235 | 109,797 | 4,048,937 | 1,522 | 4,290,491 |
| *Size-filtered ASV* | 157 | 124 | 9,636 | 26 | 9,779 |
| **Taxonomy-filtered Reads (Final)** | 78,910 | 59,696 | 4,032,426 | 1,166 | **4,172,198** |
| *Taxonomy-filtered ASVs* | 116 | 99 | 9,474 | 22 | **9,557** |
| **Percent of Raw Reads in Final Data** | 6% | 5% | 51% | 1% | **39%** |
| **Reads/Sample** |  |  |  |  |  |
| **Median** | 198 | 176 | 113,669 | 38.5 |  |
| **SD** | 1,183 | 1,332 | 53,333 | 153 |  |
| **Min** | 3 | 2 | 22,129 | 6 |  |
| **Max** | 4,206 | 5,564 | 199,759 | 442 |  |
| **Taxonomic Groups** |  |  |  |  |  |
| <u>**Metazoans**</u> |  |  |  |  |  |
| **# Metazoan Reads** | 74,952 | 53,016 | 3,321,383 | 1016 | 3,450,367 |
| **% of Total Reads** | 1.8% | 1.3% | 79.6% | 0.0% | 82.7% |
| **# ASVs** | 76 | 56 | 3,906 | 13 | 3944 |
| <u>**Non-metazoans**</u> |  |  |  |  |  |
| **# Non-metazoan Reads** | 41 | 4 | 203,015 | 64 | 203,124 |
| **% of Total Reads** | 0.0% | 0.0% | 4.9% | 0.0% | 4.9% |
| **# ASVs** | 5 | 2 | 280 | 5 | 286 |
| <u>**Unknowns**</u> |  |  |  |  |  |
| **# Unknown reads** | 3,917 | 6,676 | 508,028 | 86 | 518,707 |
| **% of Total Reads** | 0.1% | 0.2% | 12.2% | 0.0% | 12.4% |
| **# ASVs** | 35 | 41 | 5,288 | 4 | 5327 |

**Supplementary Table 3.** Individually identified and DNA barcoded cyclopoid and harpacticoid copepods, primer sets tested, sequencing result, and NCBI Accession Number if sequence was submitted to the database.

| Order | Putative Morphological ID | ID Code | Primer set | Result | NCBI Accession Number |
| --- | --- | --- | --- | --- | --- |
| Cyclopoida | Acanthocyclops americanus | CCA | mlCOLintF/<br>jgHCO2198 | good | (for our sequence) |
|  | Acanthocyclops robustus | CCB1 | LCO1490/<br>jgHCO2198 | good | N/A |
|  | Acanthocyclops robustus | CCB2 | both | no amp | N/A |
|  | Acanthocyclops robustus | CCB3 | both | no amp | N/A |
|  | Acanthocyclops robustus | CCB4 | mlCOLintF/<br>jgHCO2198 | non-target | N/A |
|  | Acanthocyclops robustus | CCB5 | both | no amp | N/A |
|  | Homocyclops spp. | CCC1 | LCO1490/<br>jgHCO2198 | good | N/A |
|  | Homocyclops spp. | CCC2 | mlCOLintF/<br>jgHCO2198 | non-target | N/A |
|  | Acanthocyclops capillatus | CCD1 | mlCOLintF/<br>jgHCO2198 | no amp | N/A |
|  | Acanthocyclops capillatus | CCD2 | mlCOLintF/<br>jgHCO2198 | no amp | N/A |
|  | Acanthocyclops capillatus | CCD3 | both | no amp | N/A |
|  | Acanthocyclops robustus | CCE1 | mlCOLintF/<br>jgHCO2198 | non-target | N/A |
|  | Acanthocyclops robustus | CCE2 | both | no amp | N/A |
|  | Acanthocyclops robustus | CCE3 | mlCOLintF/<br>jgHCO2198 | non-target | N/A |
|  | Acanthocyclops robustus | CCE4 | LCO1490/<br>jgHCO2198 | good | N/A |
|  | Acanthocyclops robustus | CCE5 | both | no amp | N/A |
|  | Mesocyclops pehpeiensis | CCF | mlCOLintF/<br>jgHCO2198 | good | (for our sequence) |
|  | Diacyclops thomasi or<br>Acanthocyclops capillatus | CCG1 | mlCOLintF/<br>jgHCO2198 | good | N/A |
|  | Diacyclops thomasi or<br>Acanthocyclops capillatus | CCG2 | mlCOLintF/<br>jgHCO2198 | non-target | N/A |
|  | Diacyclops thomasi or<br>Acanthocyclops capillatus | CCG3 | both | no amp | N/A |
|  | Diacyclops thomasi or<br>Acanthocyclops capillatus | CCG4 | mlCOLintF/<br>jgHCO2198 | non-target | N/A |
|  | Cyclops scutifer | CCH1 | mlCOLintF/<br>jgHCO2198 | non-target | N/A |
|  | Cyclops scutifer | CCH2 | both | no amp | N/A |
|  | Eucyclops elegans or agilis | CCI | mlCOLintF/<br>jgHCO2198 | non-target | N/A |
|  | Unknown cyclopoida | CCJ | both | no amp | N/A |
|  | Macrocylops distinctus | CCK | LCO1490/<br>jgHCO2198 | good | N/A |
|  | Limnoithona tetraspina | CL1 | mlCOLintF/<br>jgHCO2198 | good | (for our sequence) |
|  | Limnoithona tetraspina | CL2 | mlCOLintF/<br>jgHCO2198 | good | (for our sequence) |
|  | Limnoithona tetraspina | CL3 | mlCOLintF/<br>jgHCO2198 | good | (for our sequence) |
|  | Limnoithona tetraspina | CL4 | mlCOLintF/<br>jgHCO2198 | good | (for our sequence) |
|  | Limnoithona tetraspina | CL5 | mlCOLintF/<br>jgHCO2198 | good | (for our sequence) |
| Harpacticoida | Euterpina acutifrons | H1 | LCO1490/<br>jgHCO2198 | good | N/A |
|  | Euterpina acutifrons | H2 | LCO1490/<br>jgHCO2198 | good | N/A |
|  | Euterpina acutifrons | H3 | LCO1490/<br>jgHCO2198 | good | N/A |
|  | Euterpina acutifrons | H4 | mlCOLintF/<br>jgHCO2198 | good | N/A |
|  | Euterpina acutifrons | H5 | LCO1490/<br>jgHCO2198 | good | N/A |
|  | Euterpina acutifrons | H6 | LCO1490/<br>jgHCO2198 | good | N/A |
|  | Euterpina acutifrons | H7 | LCO1490/<br>jgHCO2198 | good | N/A |
|  | Euterpina acutifrons | H8 | both | good* | N/A |
|  | Euterpina acutifrons | H9 | LCO1490/<br>jgHCO2198 | good | N/A |
|  | Euterpina acutifrons | H10 | mlCOLintF/<br>jgHCO2198 | good | N/A |
|  | Pseudobradia sp. | HG | mlCOLintF/<br>jgHCO2198 | good | N/A |

**Supplementary Table 4.**
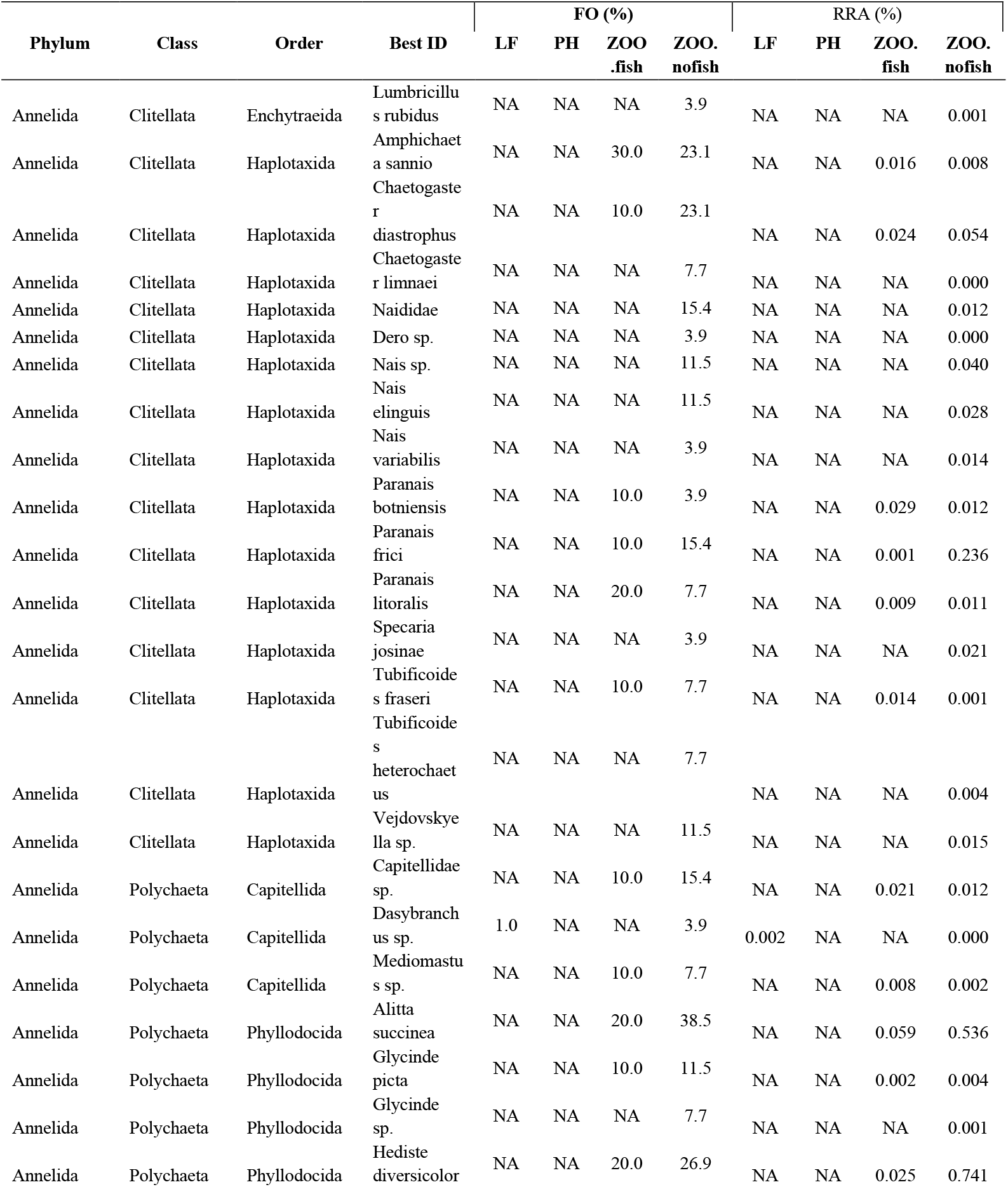
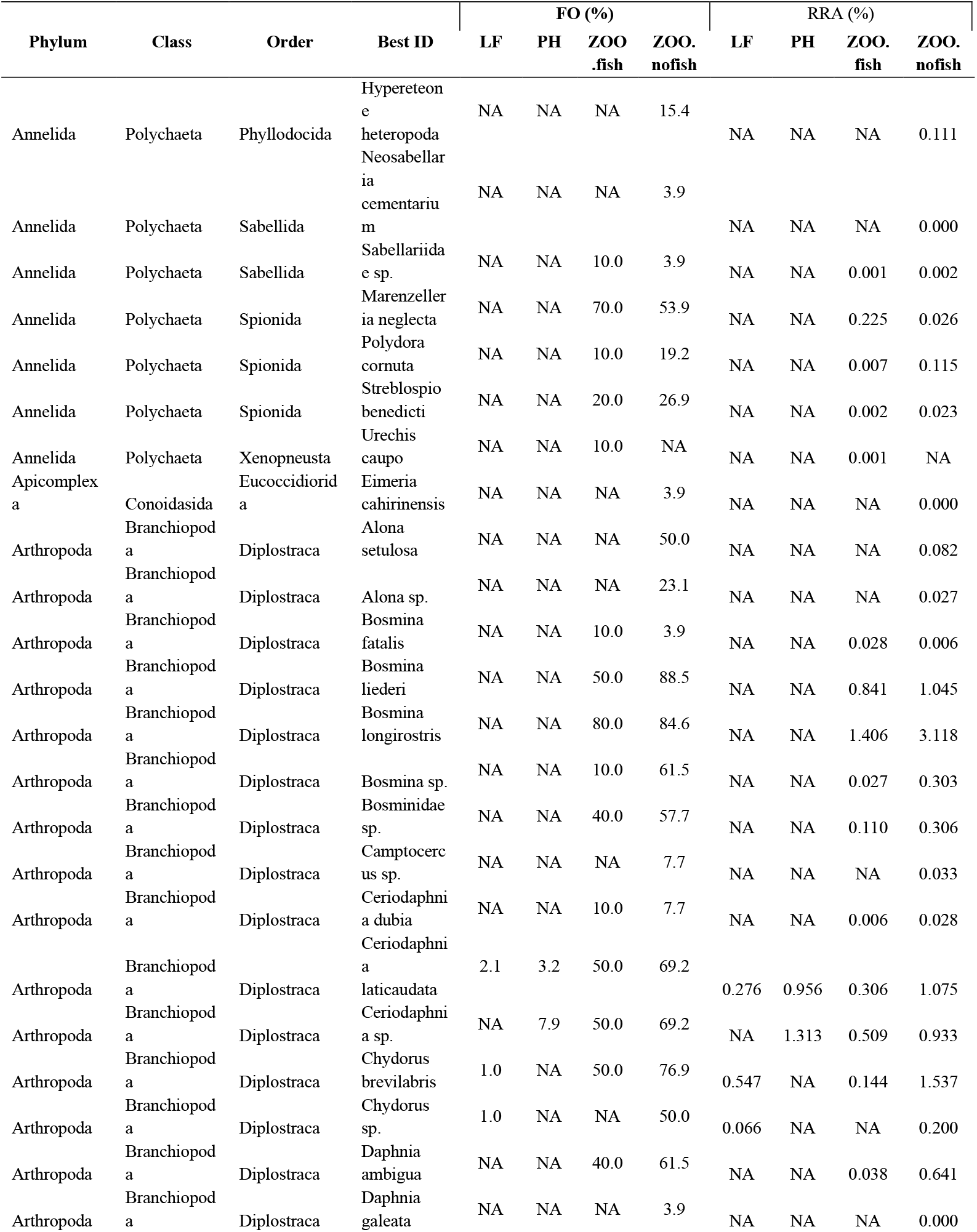
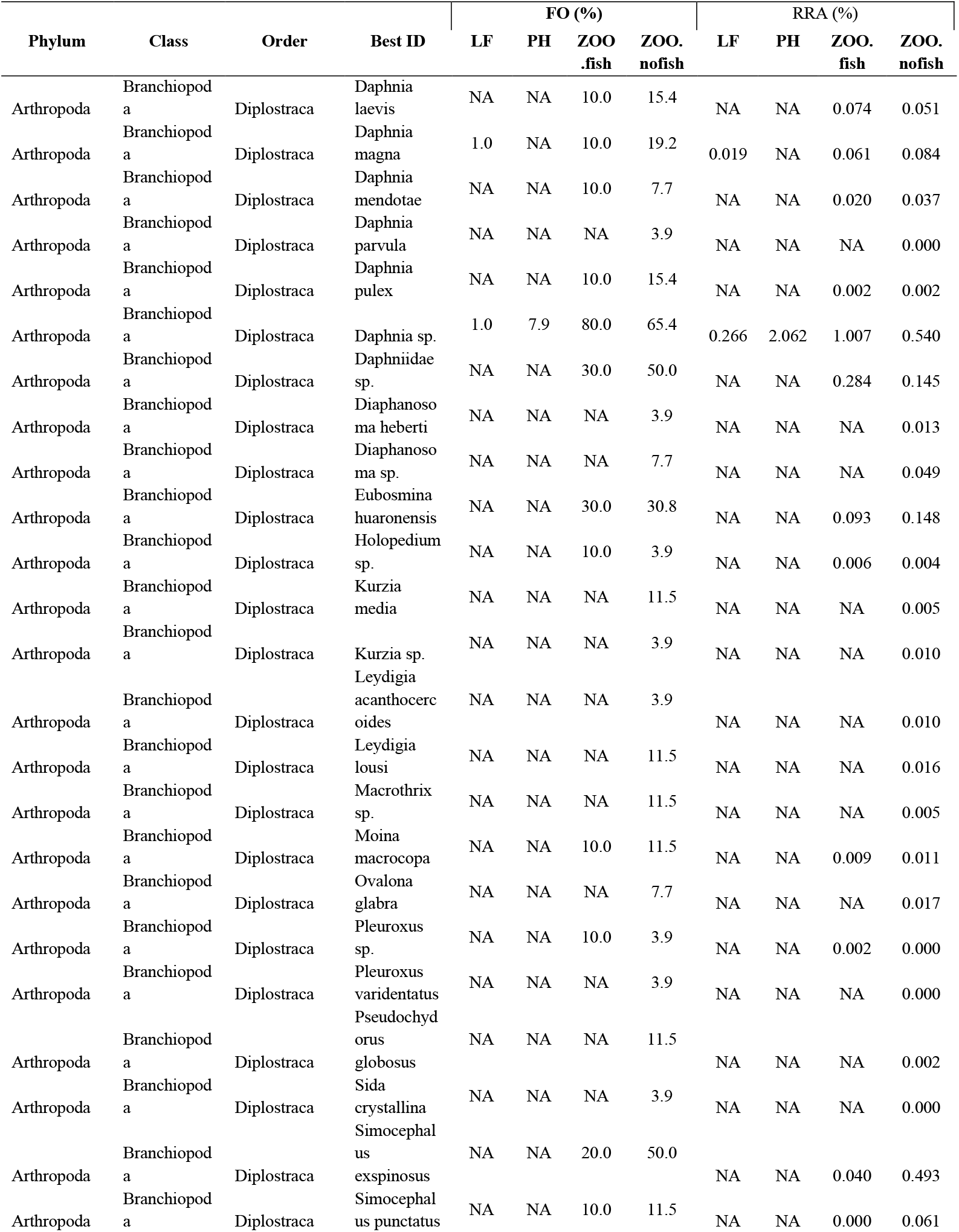
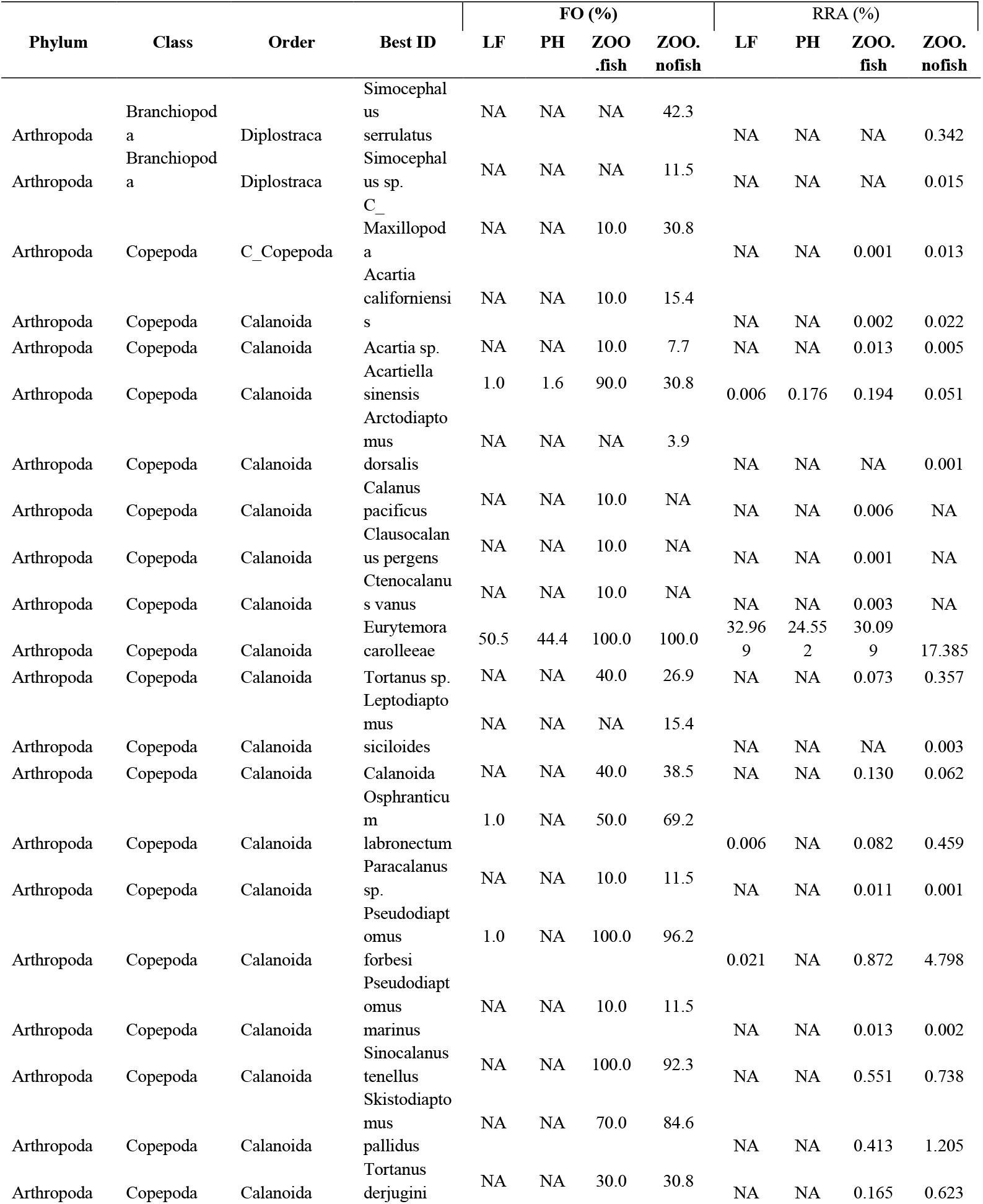
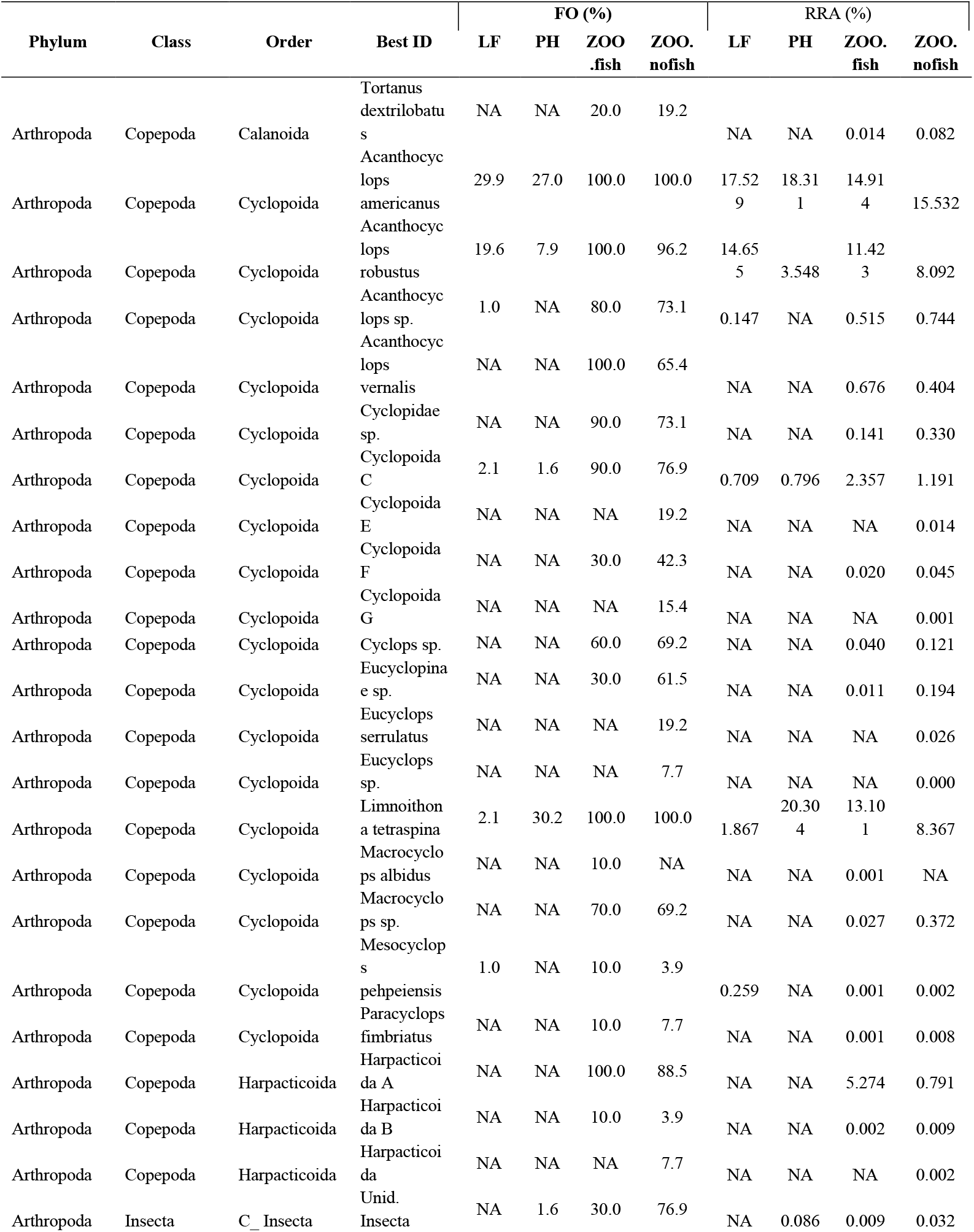
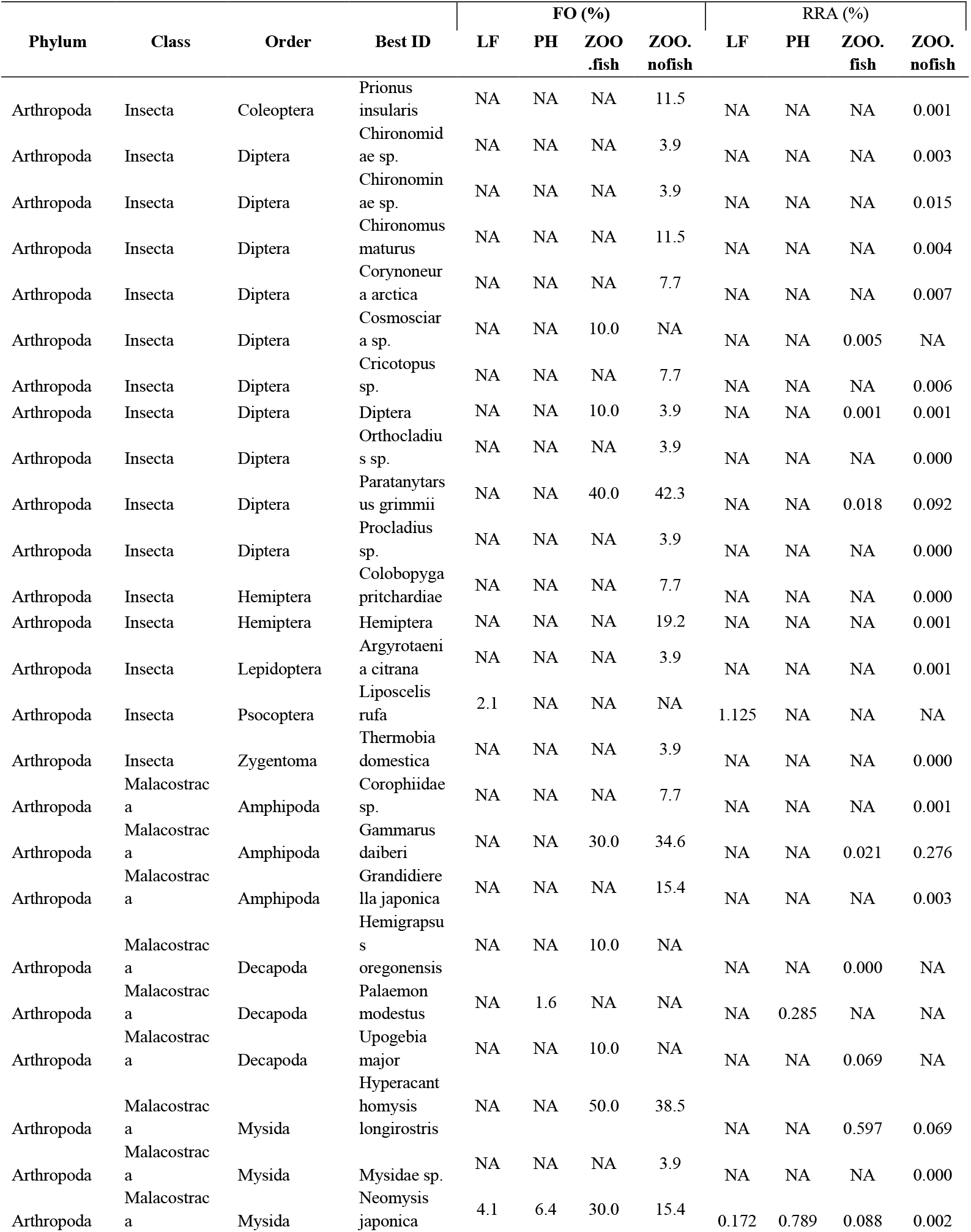
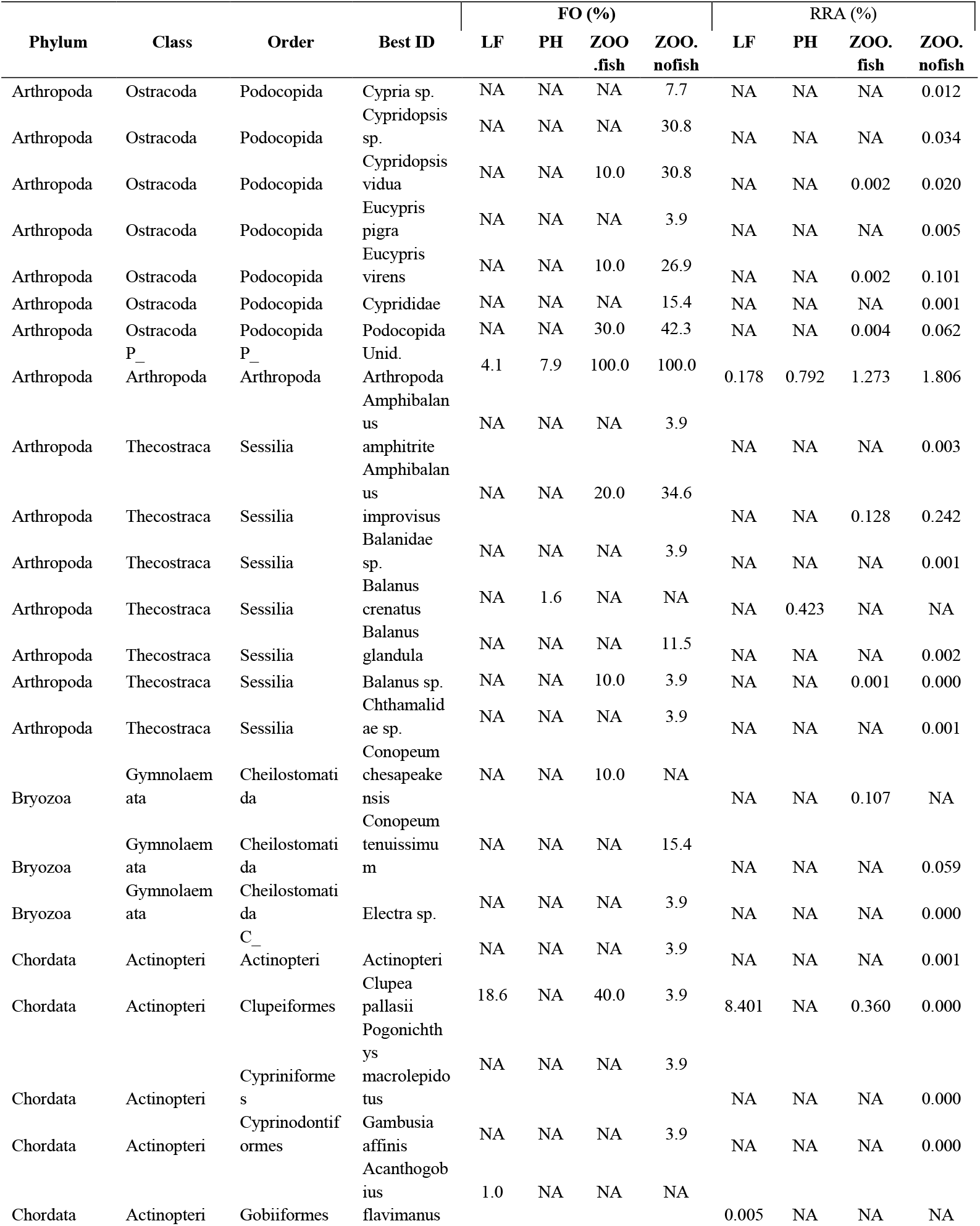
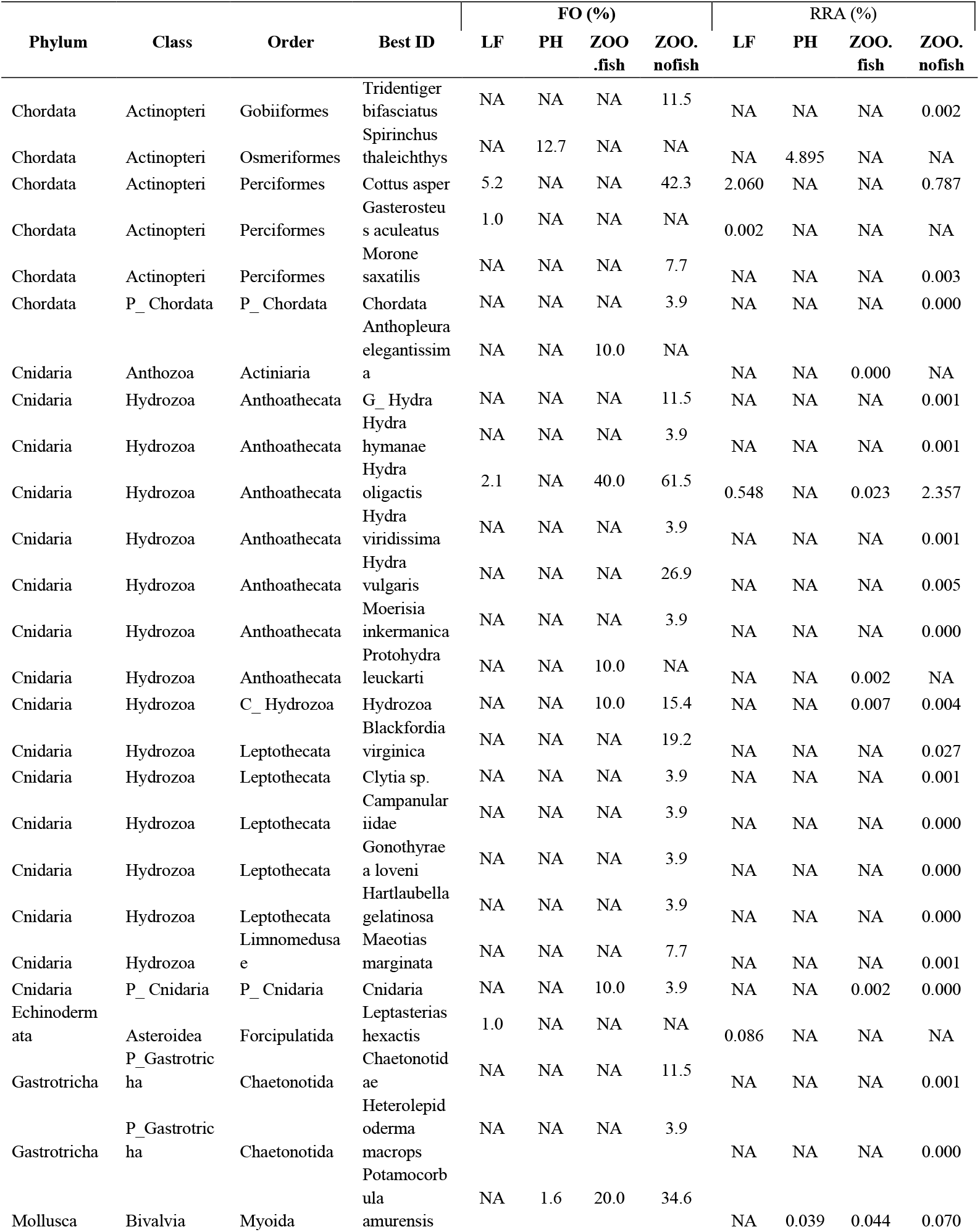
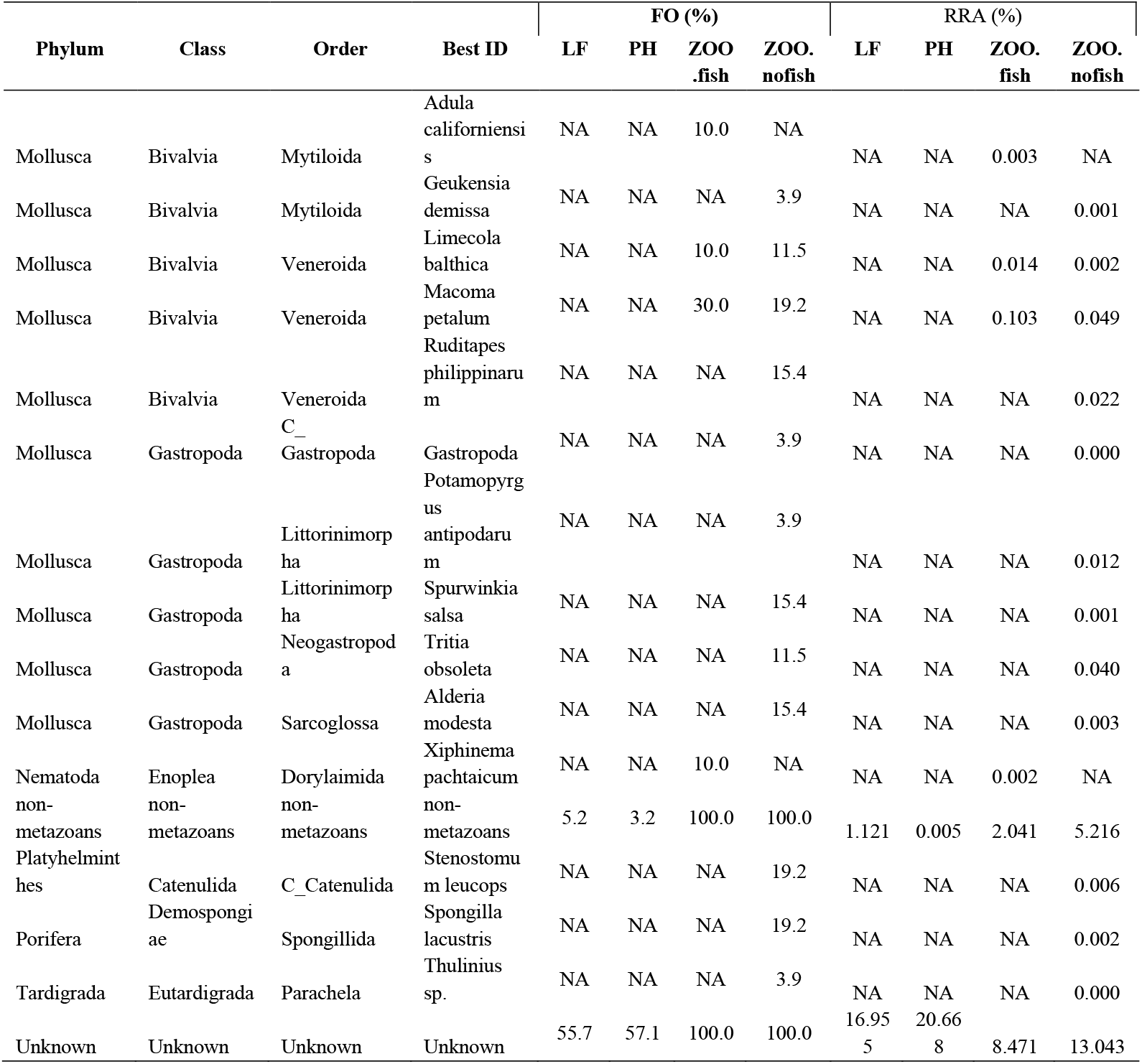
Frequency of occurrence (FO%) and Relative Read Abundance (RRA%) of high-level confidence ID’s in fish guts and zooplankton.

**Supplementary Table 5.**
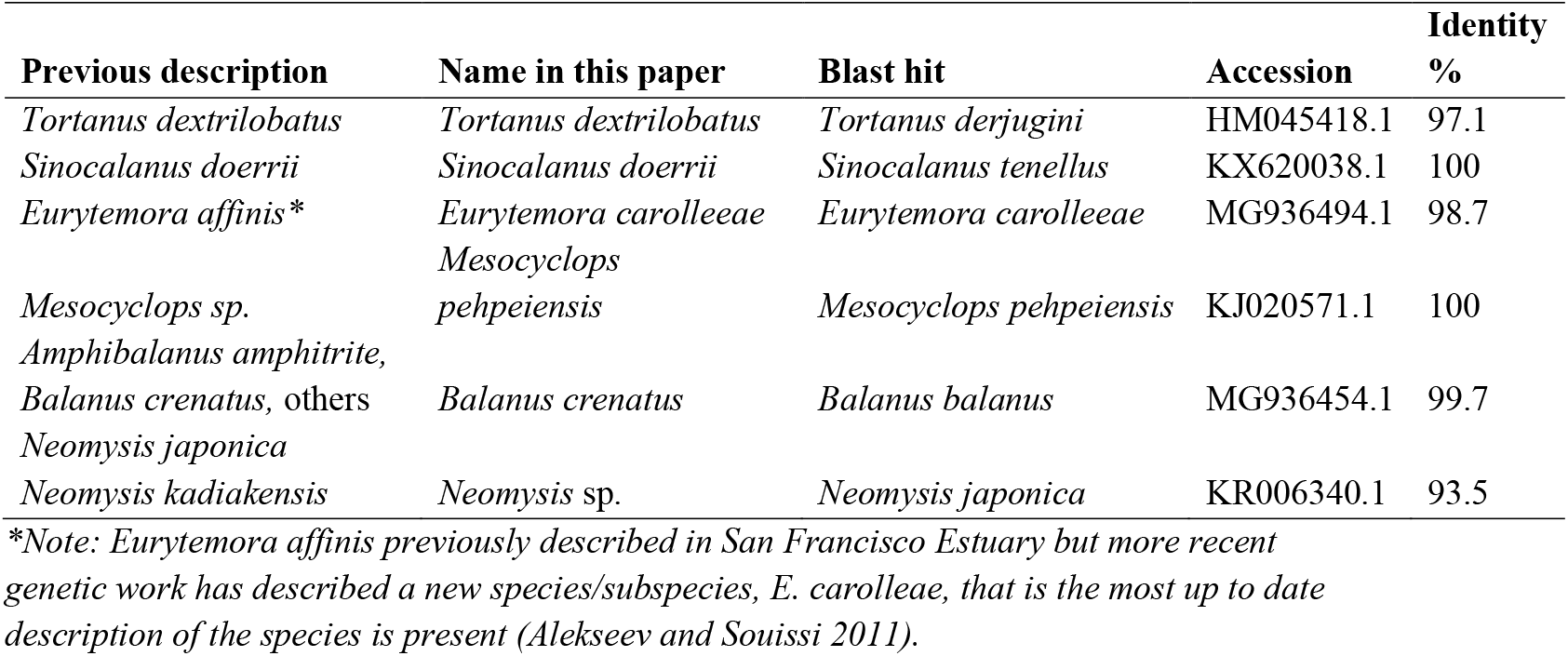
Name adjustments.

## Supplementary Figures

**Supplementary Figure 1.**
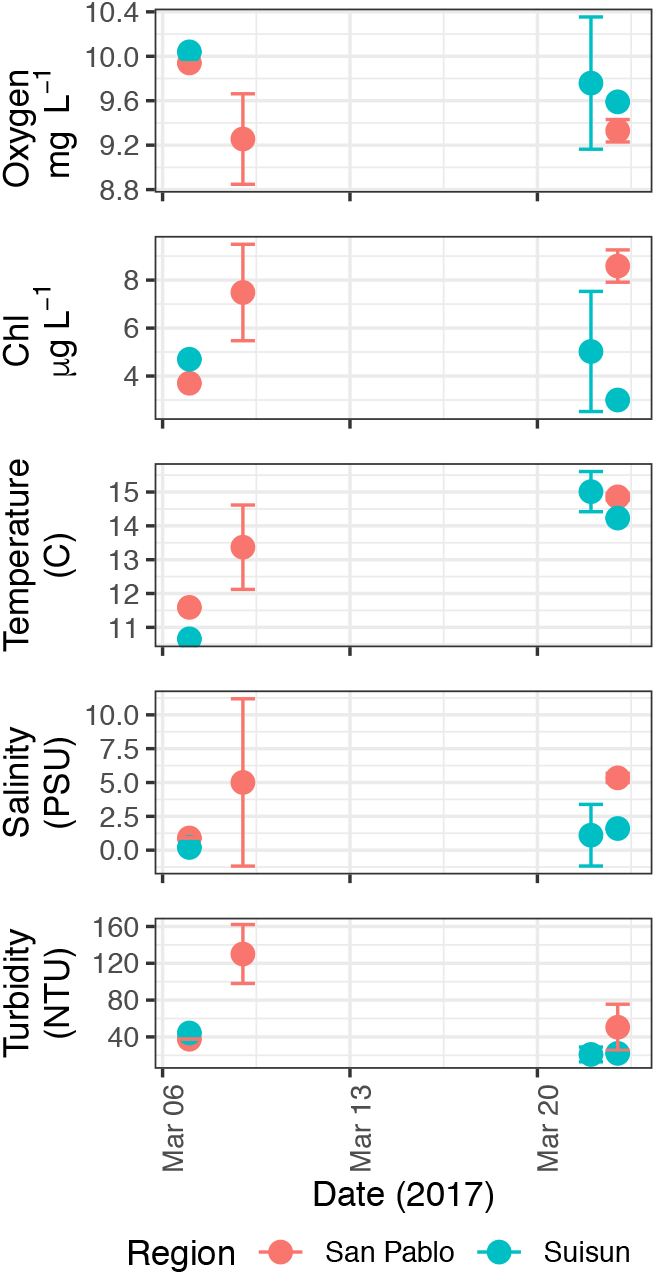
Environmental data summary for San Pablo Bay and Suisun Bay on sampling dates during this study, showing the mean and standard deviation value of each daily measurement across sites where the target species of larval fish were sampled that day.

**Supplementary Figure 2:**
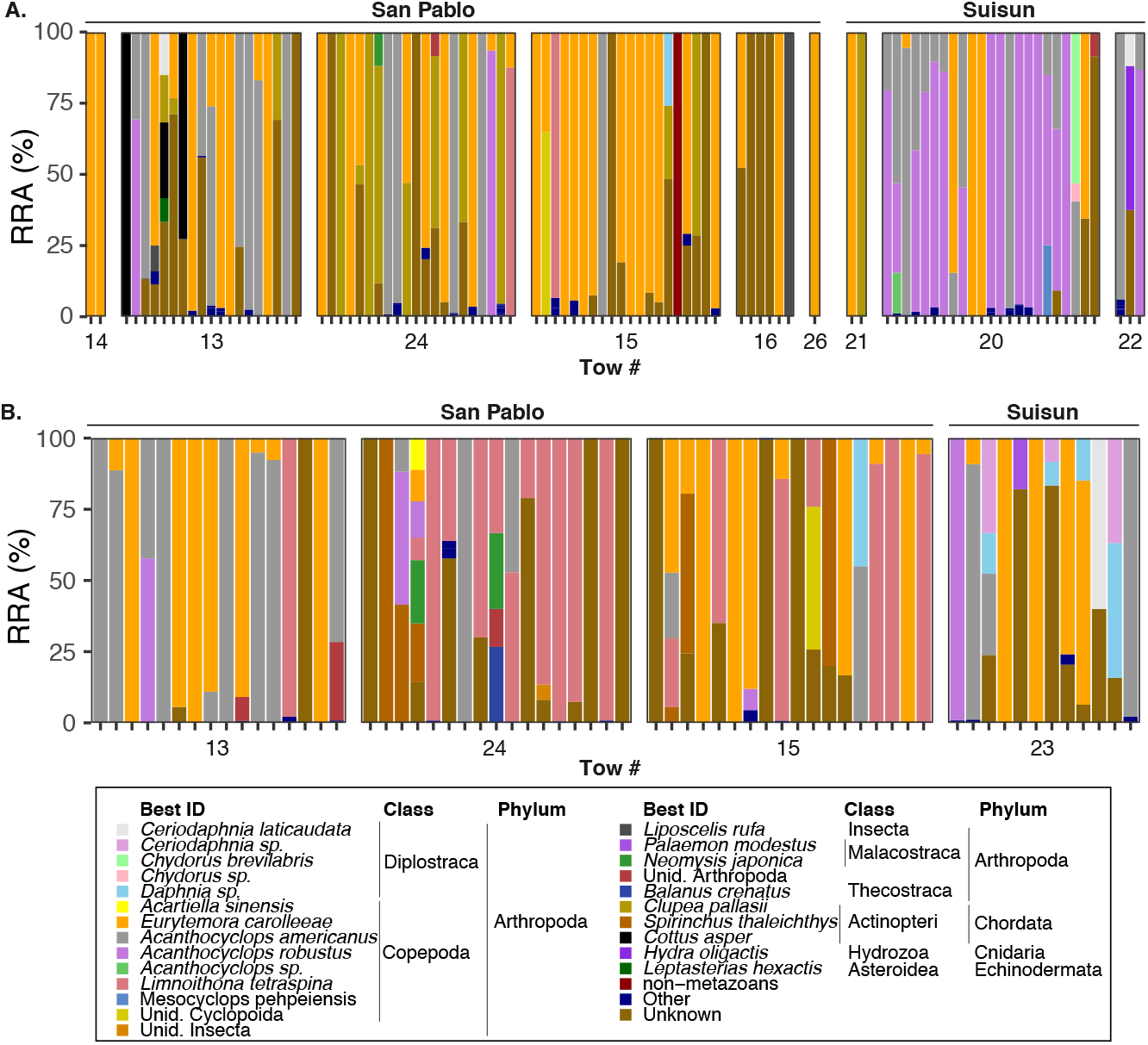
Individual variation in relative read abundances (RRA %) in fish guts for longfin smelt samples (A) and Pacific herring samples (B) across regions and tow number. “Other” group includes < 5% contribution to the sample. Unknowns are those identified with < 80% RDP bootstrap confidence.

**Supplementary Figure 3.**
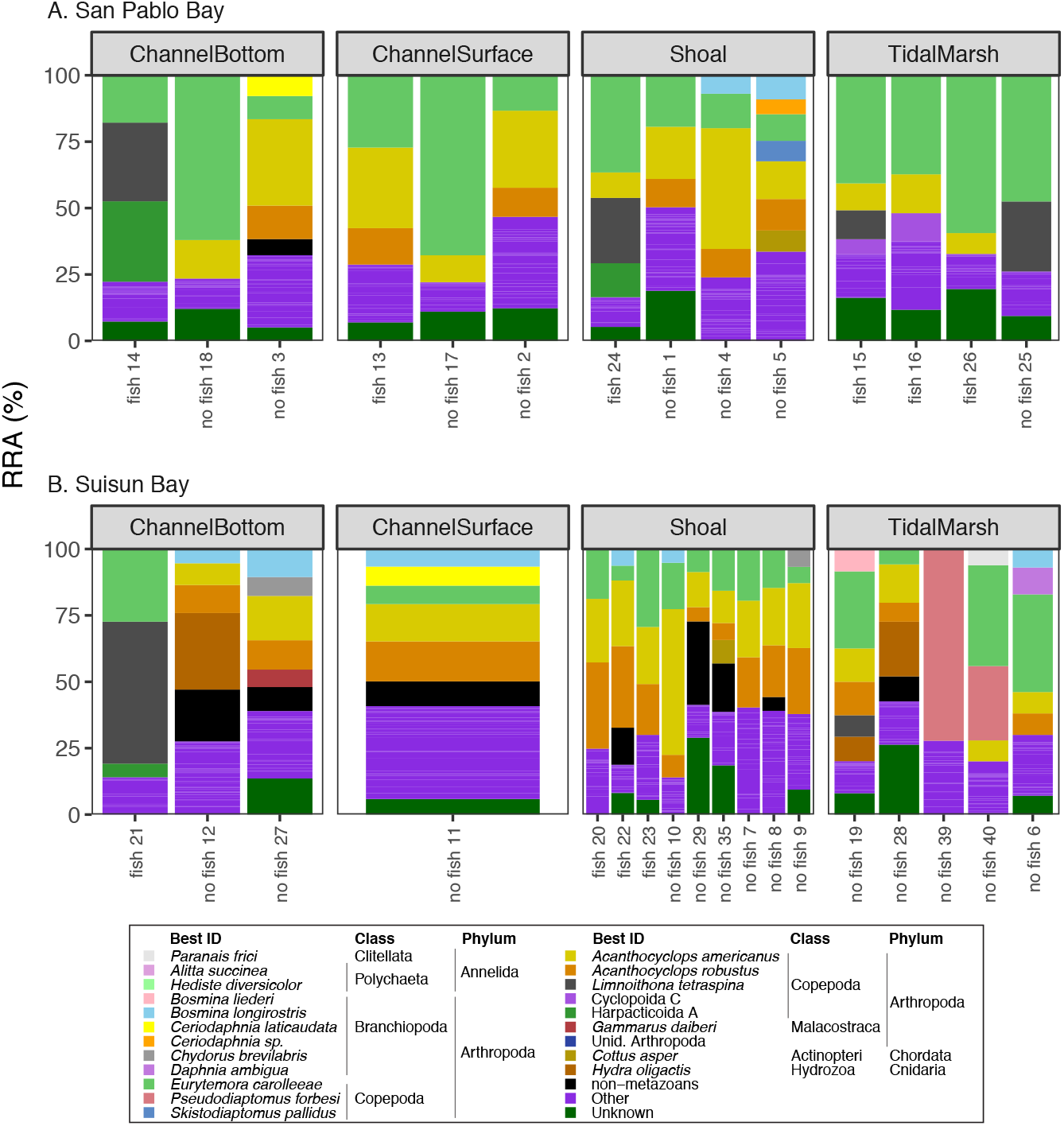
Individual variation in relative read abundances (RRA %) in zooplankton samples by habitat. Other is < 5% contribution. Tow number and whether the sample had longfin smelt concurrently collected and sequenced (fish) or not (no fish) are shown on the x-axis.

**Supplementary Figure 4.**
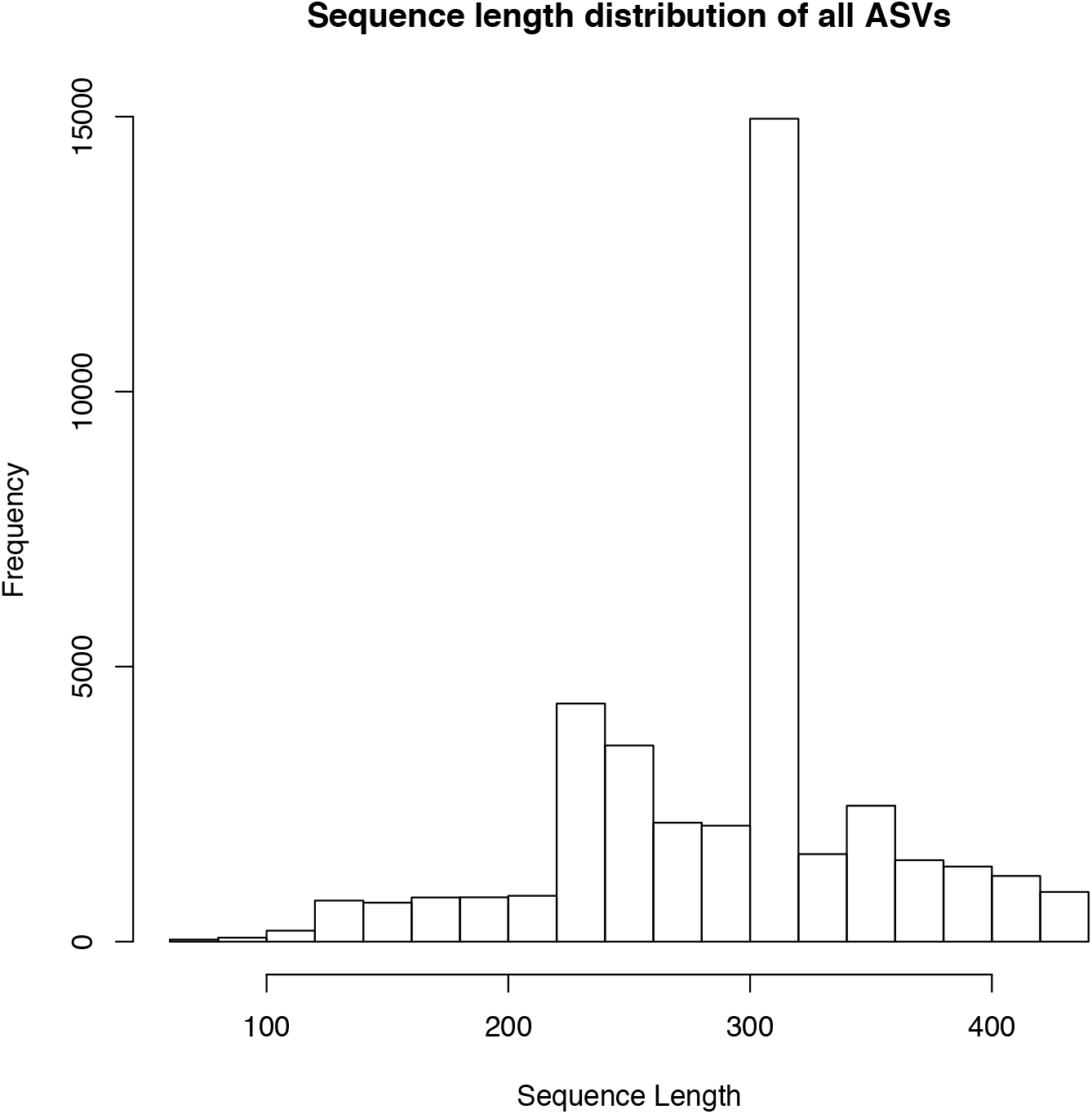
Histogram of sequence lengths in DADA2-processed set of amplicon sequence variants (ASVs).

## Notes

### Competing Interest Statement

The authors have declared no competing interest.

